# Integrated environmental and genomic analysis reveals the drivers of local adaptation in African indigenous chickens

**DOI:** 10.1101/2020.08.19.257212

**Authors:** Almas A. Gheyas, Adriana Vallejo Trujillo, Adebabay Kebede, Maria Lozano-Jaramillo, Tadelle Dessie, Jacqueline Smith, Olivier Hanotte

## Abstract

Elucidating the genetic basis of environmental adaptation in indigenous livestock populations has important implications for sustainable breeding improvement. It requires a detailed untangling of relevant environmental pressures and fine resolution detection of genomic signatures of selection associated with these environmental parameters. Here, we uniquely employed an integrative approach, combining Ecological Niche Modelling (ENM) with (i) genome-wide analyses of positive signatures of selection (SSA) and (ii) genotype-environment association (GEA) analyses to unravel the environmental adaptation of Ethiopian indigenous chickens. We first examined 34 agro-ecological and climatic variables and identified six main environmental selection drivers (one temperature - strongly correlated to elevation, three precipitation, and two soil/land variables). We then performed genomic analyses using high-density SNP data from whole-genome sequencing of 245 Ethiopian chickens from 25 different populations. Environmental genomic association analyses (SSA and GEA) identify a few strongly supported selected genomic regions, often with clusters of candidate genes, related to altitude-induced stresses (hypoxia, thrombosis, and cold temperatures), water scarcity, and the challenges of scavenging feeding behaviour. These analyses support a predominantly oligogenic control of environmental adaptation and possible regulation by one or few genetic elements only. Our study shows that the pre-identification of the key environmental drivers of adaptation, followed by a detailed genomic investigation of the associated genetic mechanisms provides a powerful new approach for elucidating the effect of natural selection in domestic animals. These results represent new landmarks to inform sustainable poultry breeding improvement.

## Introduction

A key strategy to address the health and livelihood of smallholder farming communities across developing countries is to improve their farm productivity. Ethiopia ranks first in Africa and tenth globally in terms of inventory of livestock populations (1). Here, backyard poultry farming constitutes an important economic activity, accounting for about 97% of the country’s total meat and egg production and thereby providing both income and nutrition to poor rural households and contributing significantly to the national economy. As in most of the African continent, such backyard farming still relies predominantly on indigenous poultry breeds, which are characterized by their local adaptations to various environmental challenges (e.g. extreme climatic conditions, disease, predation and scarcity of food) but lower productivity compared to commercial breeds raised under managed farming conditions (1, 2). Elucidating the genetic basis of local adaptation of these birds will have important implications for sustainable improvement of poultry production.

Chicken is an introduced species in Africa. While Egypt provides the earliest iconographic evidence of domestic chicken (3), the oldest African chicken bones were found in Ethiopia at the Mezber site (Tigray region), dated to around c.921–801 BCE (4). Molecular evidence supports at least two arrival/dispersion waves for domestic chickens in Africa (3). The first wave came likely from the Indian subcontinent around three thousand years ago, following maritime and terrestrial routes, entering Africa through today’s Egypt and the Horn of the continent. The second wave of arrival occurred during the mid-first millennium AD, along Africa’s eastern coast, following the maritime trading routes. This may have brought chicken genetic diversity from as far as Southeast and East Asia (5).

Ethiopia with its extreme altitudinal topographies, varying from below sea level to over 4,500 meters above sea level (m.a.s.l), illustrates the diversity of agro-ecologies found across tropical Africa (Figure 1A-1B). Three distinct main temperature zones are observed – cool (dega), temperate (weina dega) and hot (kolla) (6). The cool zone expands over the western and the eastern parts of the north-western Ethiopian plateau with elevation generally above 2,400 m.a.s.l., and temperature between near-freezing and 16°C. Lower elevations of the plateau (1,500 - 2,400 m.a.s.l.) constitute the temperate zone, where temperature varies between 16°C and 30°C. The hot zone is located mostly in the eastern parts of the country, where the elevation is below 1,500 m.a.s.l., and the maximum temperature can reach as high as 50°C. Across the country, precipitation varies from about 15 cm to 210 cm per annum (7). While some areas receive rainfall throughout the year, in other parts, it is mostly seasonal. Rainfall is the heaviest and most abundant in the southwest and generally decreases from South to North, mainly along the eastern lowlands (6). Different combinations of temperature and rainfall patterns have created a gradation in climatic conditions, which vary from hot-humid and hot-arid to cold-humid and cold-arid (Figure 1B). Over time, the diverse agro-climatic conditions have shaped the genetics of the chicken populations enabling them to adapt to the local agro-ecological environments and production systems (2).

**Figure 1.**
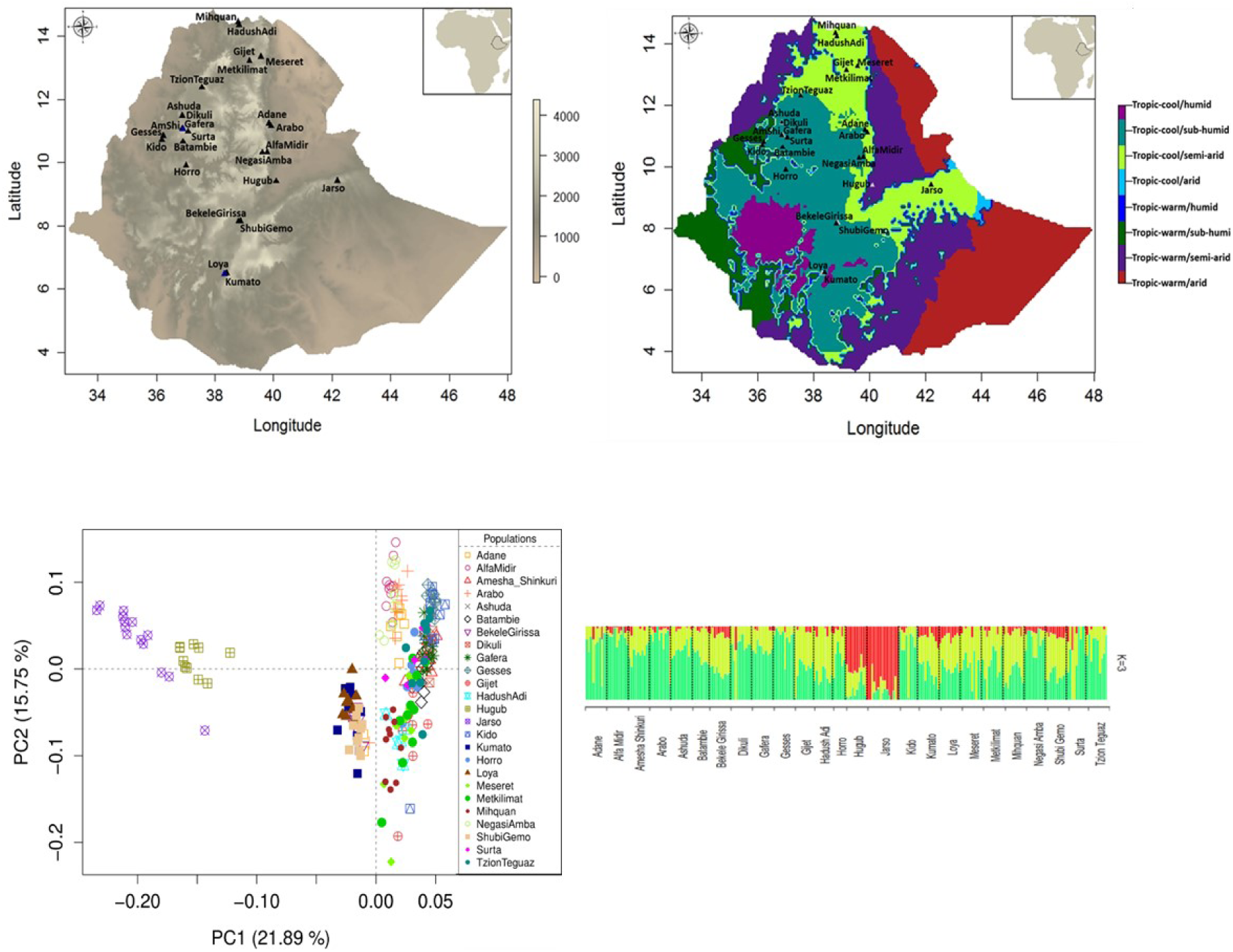
(A-B) Sampling location of Ethiopian indigenous chicken populations in relation to variation in elevation and agro-ecological zones – AEZ [62]; (C) PCA plots of the populations based on 14 million autosomal SNP; (D) Admixture analysis result with the best K value (K = 3).

Despite the observational knowledge that African indigenous chickens are adapted to their harsh environmental conditions, the genetic mechanisms underlying these adaptations are still largely unknown (8). Likewise, the environmental factors that trigger adaptive response have not been dissected thoroughly (8). Conventionally, environment-genome adaptation studies have focused on the adaptation to an inferred specific environmental stressor, e.g. high altitude or heat stress (9, 10), or in a single ecotype (11, 12), without analysing the environmental stressors of the considered agro-ecology. In the present study, we are now adopting a powerful integrative approach - combining Ecological Niche Modelling (ENM) with genomic analyses (selection signature and genotype-environment association) to first dissect the environmental drivers of local adaptation and then to investigate their impact on the genome. We apply this approach across ecotypes of Ethiopian indigenous chicken populations.

## Results

### Genomic diversity of Ethiopian indigenous chickens

Genomic data for the present study originated from the whole genome sequencing (WGS) of 245 Ethiopian indigenous chicken samples from 25 different populations representing diverse agro-climatic conditions (Figures 1A-1B, Table S1). Analysis of the WGS data detected 19.5 M SNPs, of which around 29% are novel. The genetic diversity of the populations is similar, with 10 – 12 M SNPs detected per population and a mean genome nucleotide diversity (*π*, based on individual sites) between 0.28 and 0.34. After applying stringent quality filtration, we used 14 M autosomal SNPs and 238 individuals for all downstream genomic analyses (see Methods).

Principal component analysis (PCA) based on the filtered variants reveals the structure and relatedness of the 25 populations (Figure 1C). Only the Hugub and Jarso populations from the Rift Valley slope in eastern Ethiopia are clearly separated from the other populations whereas only a weak sub-structuring based on geographic closeness is generally observed among the rest of the populations. Admixture analysis conforms to this result by showing contributions from three ancestral gene pools, with Hugub and Jarso having a major contribution from a single gene pool, which has a minor presence in the other populations (Figure 1D).

### ENM reveals the environmental diversity of Ethiopian chicken habitats and identifies important environmental drivers of local adaptation

ENM is a powerful tool for predicting the distribution of a species based on the environmental conditions of the species’ known occurrence locations. The distribution models were built using the maximum entropy algorithm implemented by MaxEnt (13) with data on 34 different agro-climatic variables at 250 geographic data points (10 per population). These were considered as “occurrence data”. Moreover, 10,000 geographic data points from the remaining of Ethiopia were included as background points, against which the occurrence data could be projected to create population-specific environmental suitability maps. The agro-climatic variables included 21 climatic parameters, eight soil properties, four vegetation parameters, and elevation data from public databases (Table S2). These variables were chosen considering their biological relevance for scavenging chickens, e.g. climatic variables and elevation are expected to affect physiological tolerance of chicken, soil variables may influence the type and abundance of food, and the vegetation parameters may affect both food availability and exposure to predation. Accordingly, these variables were considered as proxies of environmental selection pressures.

In the first step of ENM, we removed variables which are highly correlated (*r*_*s*_ > 0.6; except one from each correlated group) and/or with low contribution (< 4%) (Figures S1, S2). It retained only eight variables: the minimum temperature of the coldest month of a year (minTemp), precipitation seasonality which represents the variation in precipitation across a year (precSeasonality), precipitation in the wettest quarter (precWQ) of a year, precipitation in the driest quarter of a year (precDQ), soil organic carbon content (SoilOrgC), grass/shrub cover (Grassland), proportion of cultivated land (LandUse) and the dominant cultivated crop in an area (Crop_dominance). Upon further checking, the Grassland variable was removed as it showed high multi-collinearity (Variance Inflation factor > 7) with LandUse. We also removed Crop_dominance because of ambiguity and possible erroneous categorization of some of the data points based on visual examination in Google Earth.

The final model with the six selected variables produced a refined estimate of the relative contribution of the variables (Figure 2A). SoilOrgC shows the largest individual contribution (24%), followed by the minTemp (21%), while LandUse has the smallest contribution (10%). The three precipitation variables (precWQ, precDQ and precSeasonality) show a combined contribution of 45%. PCA based on these six variables spread the populations in the environmental space, showing large heterogeneity in Ethiopian agro-climatic conditions and supporting the importance of these variables as environmental drivers of adaptation (Figure 2B). Geographically close populations are generally positioned close to each other with some notable exceptions, e.g. Alfa Midir/Negasi Amba and Arabo/Adane which are geographically close to each other but distant in the environmental space. In contrast, Arabo and Jarso appear close to each other in the environmental space even though they are geographically distant (409 km; Figures 1 and 2B). These outlier pairs show the drastic change in Ethiopian climate and landscape even within short geographic distances. The environmental diversity of Ethiopia is further illustrated by the environmental suitability maps (Figure 2C), which describe how similar is the environment across the country for each of the sampled populations.

**Figure 2.**
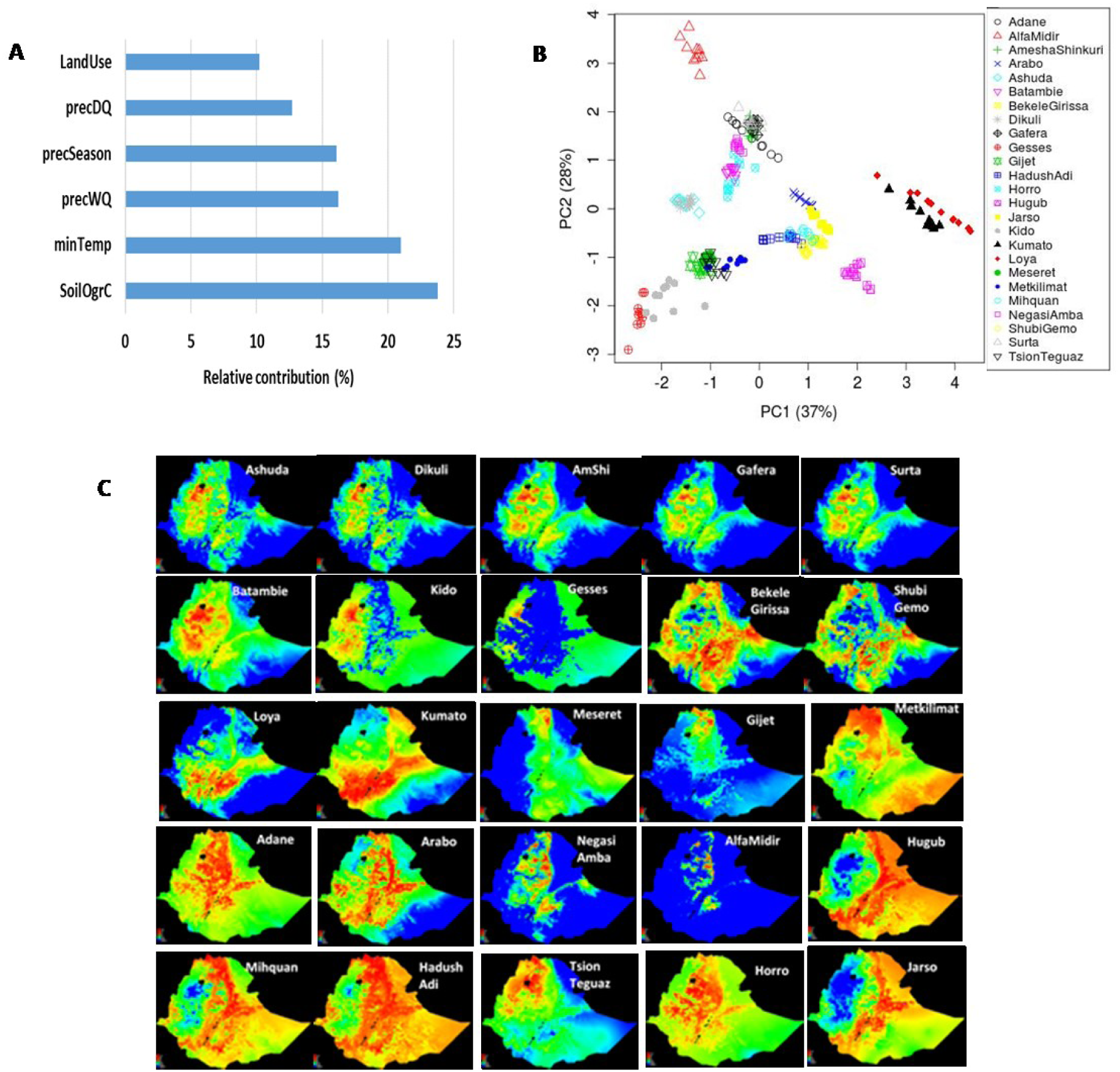
(A) Relative contribution of the six environmental variables selected based on Ecological Niche Modelling; (B) PCA plots showing the distribution of the 25 Ethiopian chicken populations in the environmental space provided by the six selected environmental parameters; (C) Suitability maps of the 25 Ethiopian chicken populations produced by ENM using six selected environmental variables. Hotter colours (towards red spectrum) indicate more suitable conditions.

### Genomic analyses identify candidate loci for environmental adaptation

With the identification of the important environmental drivers of selection, our next goal was to determine the genetic basis of adaptation to these factors. Two types of analyses were performed: (i) Selection Signature Analysis (SSA) by comparing extreme groups of populations (Low *versus* High) for each environmental predictor (Figures 3A, S5-S10), using *Fst* (14) and XPEHH (15) approaches with overlapping sliding windows (20 kb size with 10 Kb step) and (ii) Genotype-Environment Association (GEA) using Redundancy Analysis (RDA), a multivariate linear regression approach that can simultaneously analyze many loci to detect weak multi-locus signatures of selection (25). RDA was chosen over other GEA methods for its robust performance across different sample sizes, levels of population structure, and demographic histories (25).

**Figure 3.**
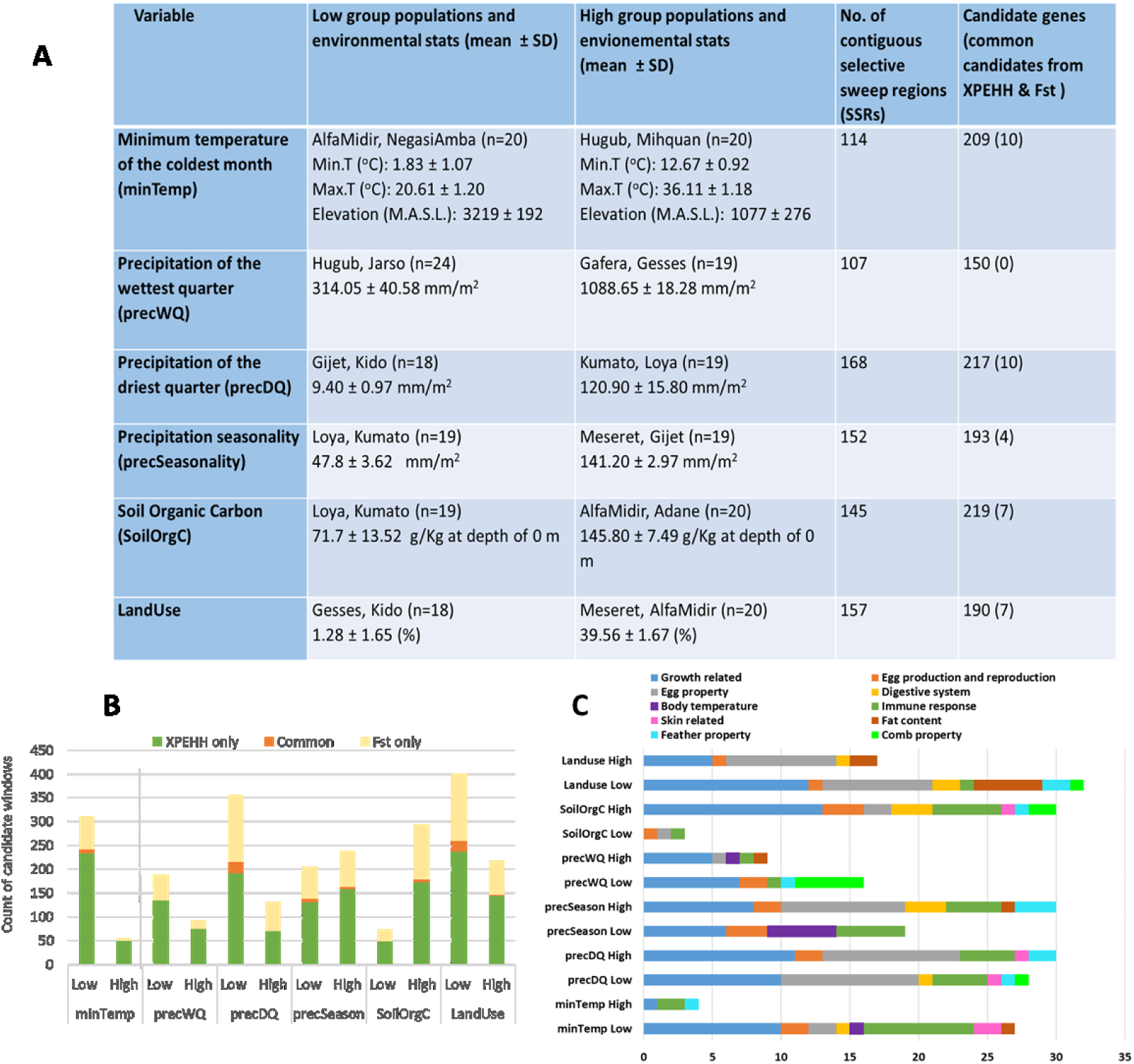
**(A)** Summary table describing the Low and High groups and selection signatures results from different environmental analyses; (B) Stacked bar plot showing the split of candidate sweep windows based on Low/High groups and detection methods; (C) Overlap of candidate genes with known QTLs from chicken QTLdb.

SSA windows with empirical *P-*value < 0.01 were considered as putative selective sweeps for a standardized *Fst* (*ZFst*) > 5 or an absolute standardized XPEHH (|XPEHH_std|) > 3 (Figures S12-S13). Moreover, since the positive and negative values of XPEHH indicate directionality of selection, all SNPs within a XPEHH-based candidate window needed to show the same directionality. For *Fst*-based candidate windows, we determined the direction of selection based on which group (Low or High) had the lower value of pooled heterozygosity (*Hp*) (16), and/or based on the signs of the XPEHH value for the corresponding windows (see Methods).

Across the different environmental analysis, we observe a weak positive correlation between the *Fst* and the XPEHH results (*r*_*s*_= 0.22 - 0.34, *P* < 2.2e-16). A similar observation has also been reported in previous studies (17). Depending on the environmental variables, *Fst* identified 71 to 237 and XPEHH 210 to 405 windows above the assigned thresholds (Figure 3B). Selective sweep windows common to both analyses were considered as our strongest candidates. Their number ranges from 6 (SoilOrgC variable) to 24 (precDQ and LandUse variables) (Figure 3B), with no window in common between the *Fst* and the XPEHH results for the precWQ variable. After merging adjacent windows 107 to 168 Selective Sweep Regions (SSRs) of 20 kb to 550 kb size range were obtained (Figure 3A). Around 76% - 90% of the detected SSRs overlap with genes (Table S3). The SSRs include a large number of SNPs (∼62,000 to ∼101,000), but only 1.5% to 8% show a large difference in allele frequency (dAAF > 0.5), and only a handful of these (n = 1 to 35) belongs to a potentially functional category (non-synonymous, splicing and ncRNA exonic) (Table S4). Hypergeometric tests show that intergenic SNPs are over-represented among those with large dAAF, while SNPs within genes (non-synonymous, intronic and UTR) are under-represented (*P* < 7.34e-09). This result is predictable, as the majority of the intergenic variants are expected to be potentially neutral, and hence these can be hitchhiked easily to high frequency with causal variants without having any physiological consequence. Many of the SSRs overlap with known QTLs (18), suggesting the affected phenotypes (Figure 3C, Table S5).

RDA was performed using a set of genome-wide LD-pruned SNPs (n = 1,210,311) as response variables and the six environmental predictors as explanatory variables. Conditioning on geography (latitude and longitude) was applied to correct for spatial autocorrelation and neutral population genetic structure arising from geographic proximity (19). The overall model was highly significant (permutation analysis *P*-value < 0.01), although it explained only 1.9% of the total genetic variance. This result is not unusual given that only a small proportion of the 1.2 M SNPs is expected to be associated with environmental predictors. The first five of the six RDA axes were significant, explaining about 91% of the variance captured by the RDA model (Figure 4). Therefore, SNPs from the two extreme ends of the loading distribution (SD > 3.5) at each significant axis were taken as outliers (n = 2,240 in total). Using the strongest correlated environmental variable for each outlier resulted in 235-571 candidate SNPs per environmental predictor. The correlation values were generally low to moderate ranging from 0.05 to 0.50 (median = 0.22) (Figure 4C). Since an LD-pruned SNP set was used in the analysis, it is quite possible that the actual functional variants were not included. We, therefore, identified any variants that are in high LD with the outliers SNPs (*r*^*2*^ > 0.8). It added 30 SNPs to the candidate SNP list, taking the total to 2,270 (Table S6). The RDA candidate SNPs represent 1,879 gene-environment combinations (212 - 446 genes per environmental analysis; Figure 4D). Only 2.8% of these are common with those detected in the SSRs (*Fst* and/or XPEHH results; Figure 4E).

**Figure 4:**
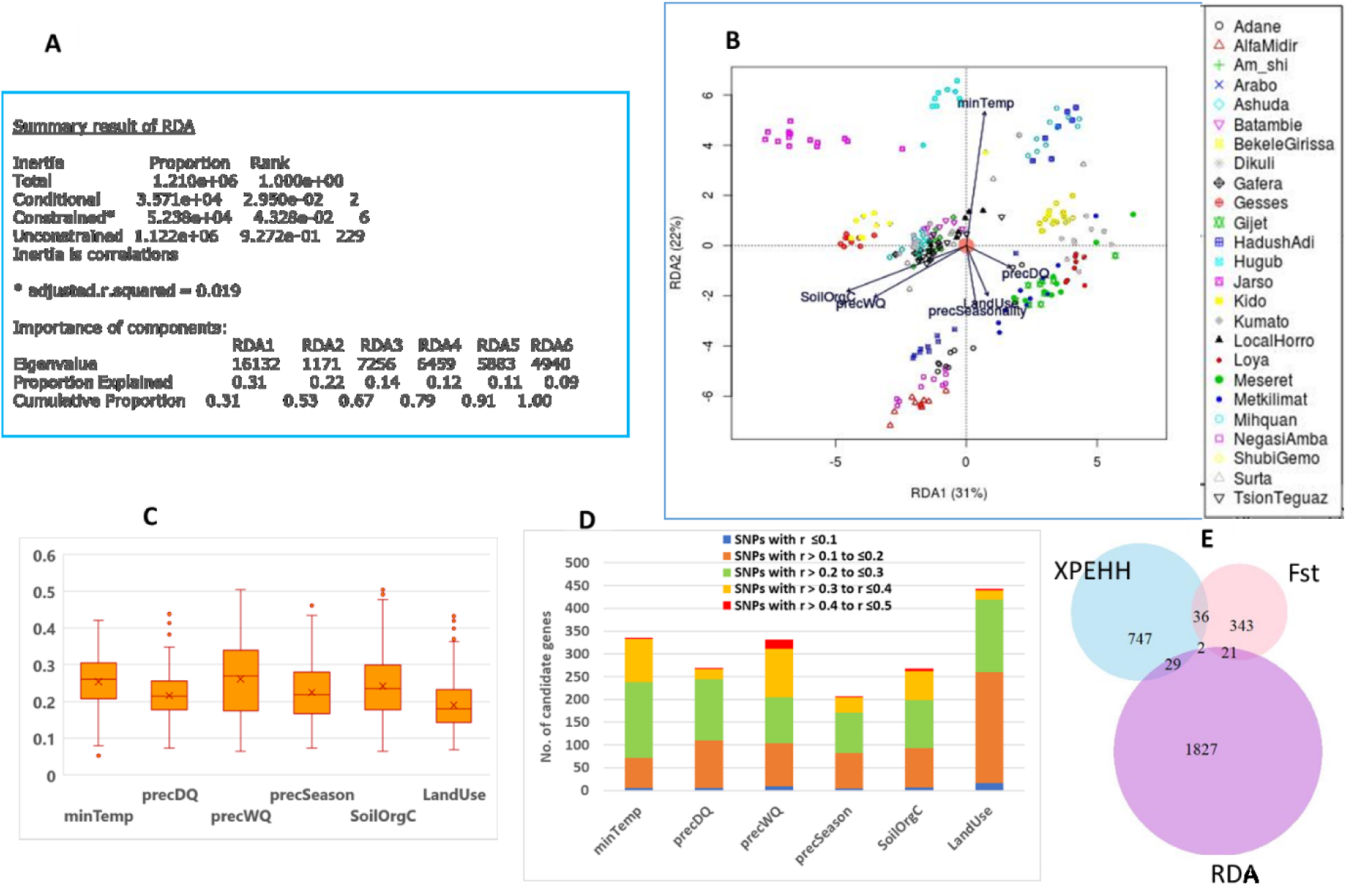
(A) Summary of Redundancy Analysis (RDA). (B) PCA plot based on RDA axes 1 and 2; (C) Box plots showing the distribution of correlation values of outlier SNPs associated with different environmental predictors; (D) Stacked bar graph showing the number of candidate genes per environmental predictor and their split based on environmental correlation; (E) Venn diagram showing overlaps of candidate genes between selection signatures and RDA analyses.

### Adaptation to extreme temperatures and high altitude

The minimum temperature of the coldest month of a year (minTemp) shows a strong positive correlation with the maximum temperature of the warmest month (*r*_*s*_ = 0.9) and a strong negative correlation (*r*_*s*_ < -0.91) with elevation (m.a.s.l.). Therefore, for the SSA, the Low group included two populations (Alfa Midir and Negasi Amba) living at the lowest minimum temperature, the lowest maximum temperature and at the highest altitude environments, while the High group included two populations (Hugub and Mihquan) living at the highest minimum temperature, highest maximum temperature and the lower-altitude environments (Figure S5). The majority of the SSRs (82%) and the strongest signals were detected in the Low group providing evidence in support of adaptation to low temperature and/or high altitude (Figures 3B, 5B-5D, Table S3). Of the 209 genes overlapping the SSRs, ten genes - all in the Low group - were detected in both *Fst* and XPEHH analyses. These are considered as the most significant candidates (Table 1). Nine of these genes (*CLP1, YPEL4, ENSGALG00000007381, UBEL6, TIMM10, RTN4RL2, SLC43A3, PGR2/3, P2RX3)* belong to a single SSR on chr5:17250000-17280000 (Figure 5G), while the remaining one (*UTP18)* overlaps the region on chr18:5100000-5120000.

**Table 1:**
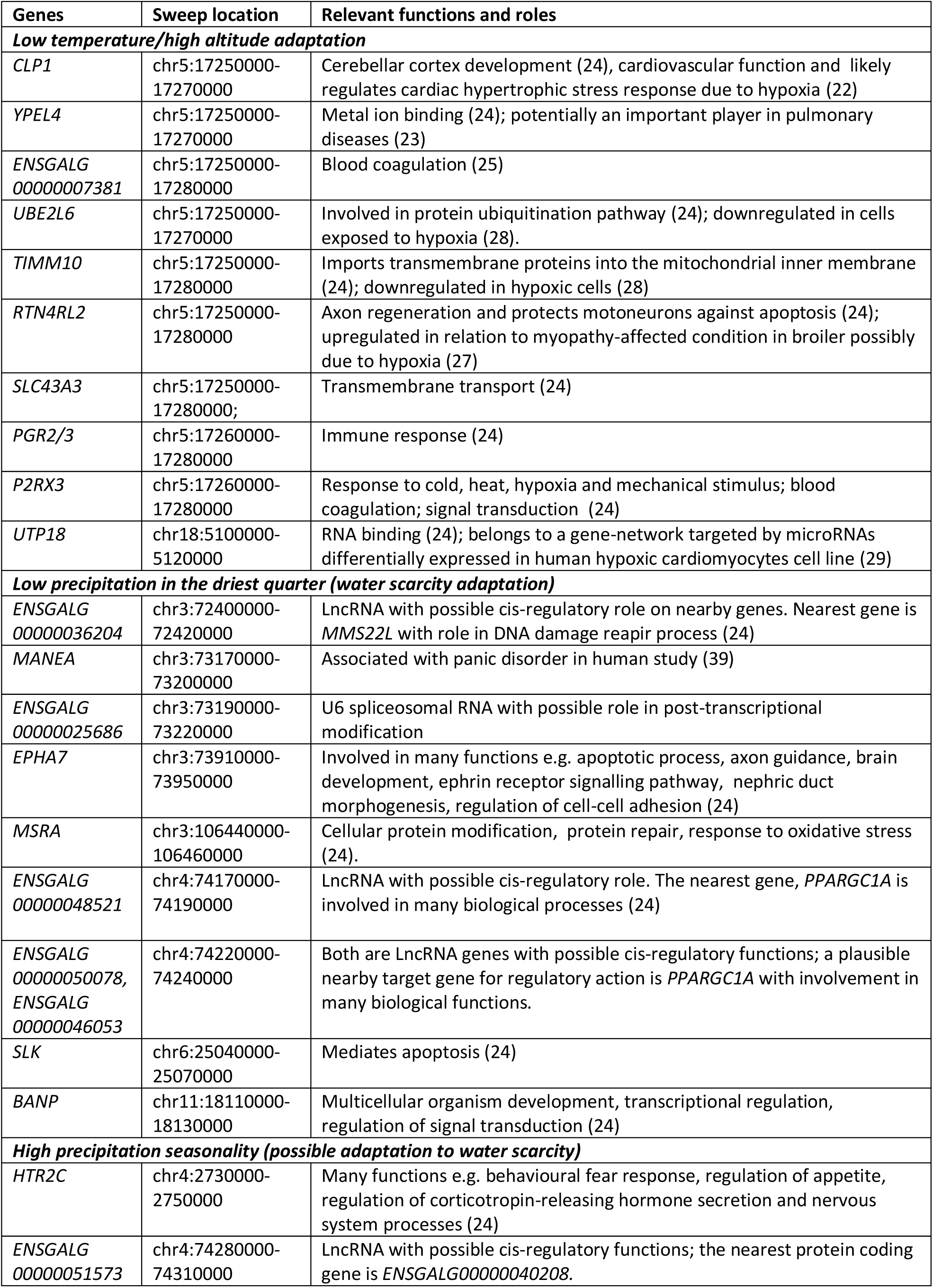
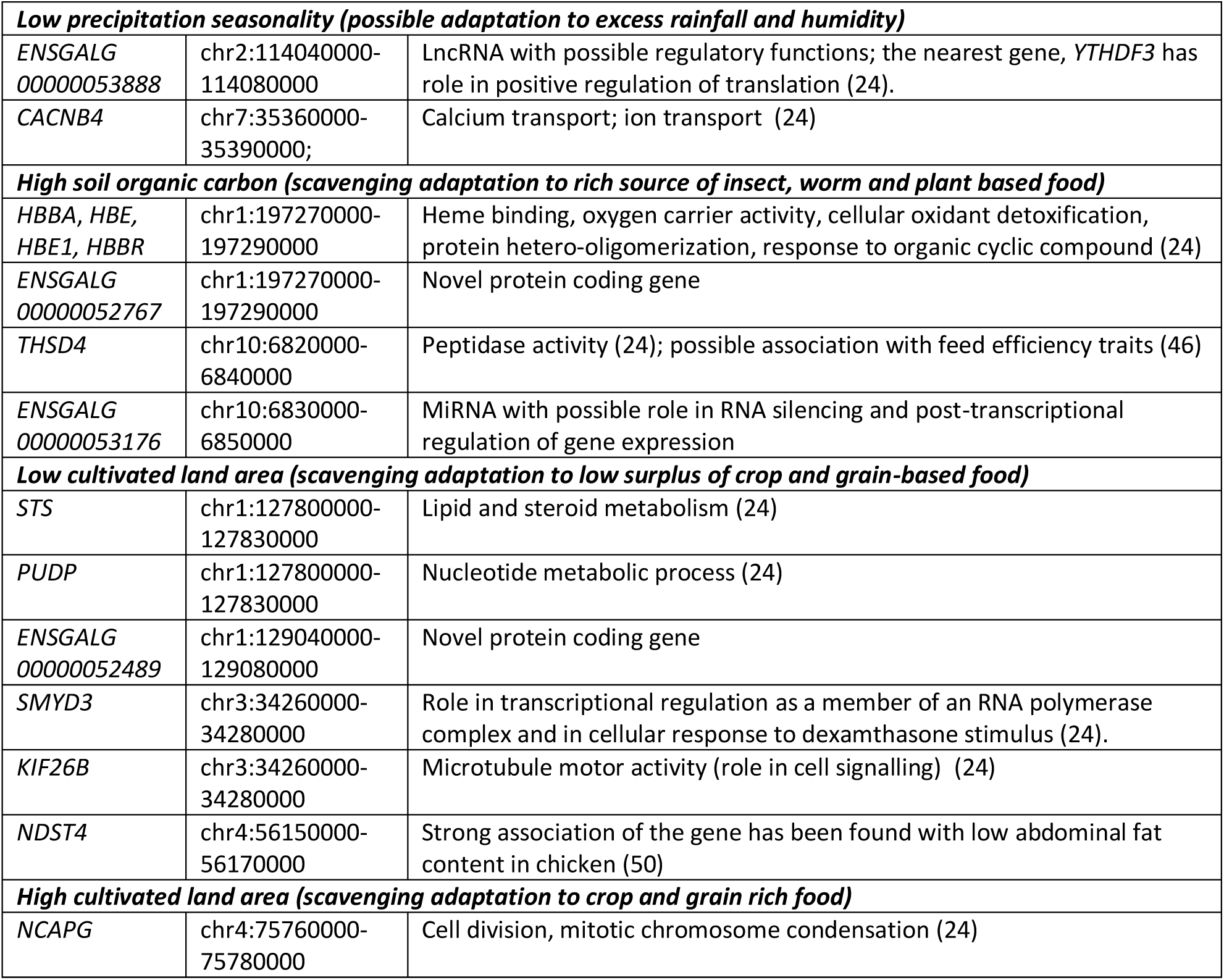
Candidate genes detected by both XPEHH and *Fst*.

**Figure 5:**
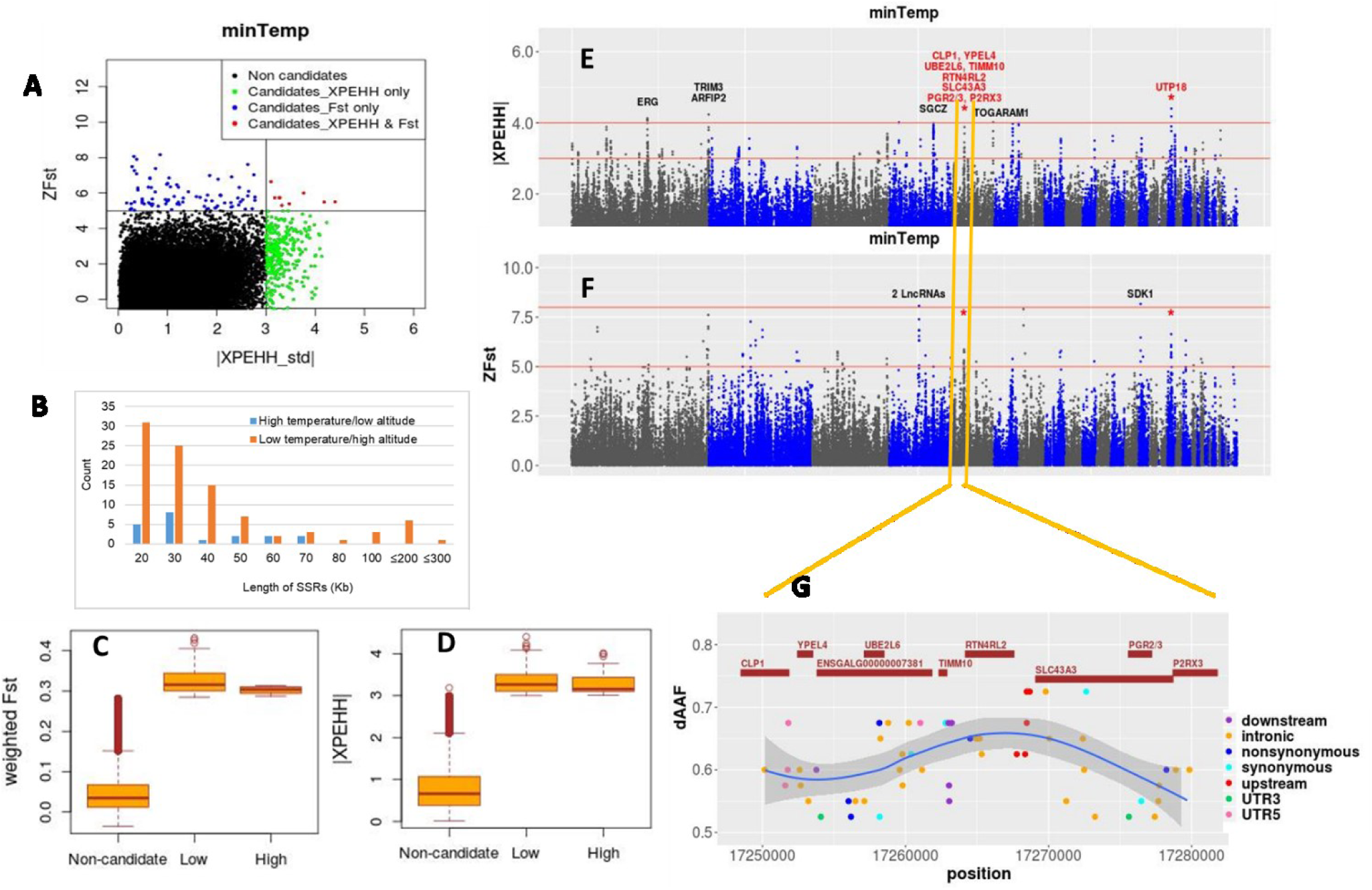
Selection signature analysis results for minTemp. (A) Scatter plot of standardized values of XPEHH versus *Fst*. (B) Length distribution of Selective Sweep Regions (SSRs). (C-D) Box plots showing the distribution of *Fst* and XPEHH metrics for non-candidate and candidate windows. (E-F) Manhattan plots for the XPEHH and *Fst* analyses; common windows are marked with asterisk and gene names from common windows are shown in red. (G) Closer look of the common *Fst*/XPEHH region - chr5: 17250000-17280000 - with SNPs showing allele frequency difference (dAAF) > 0.5 between the Low (AlfaMidir, NegasiAmba) and High (Hugub, Mihquan) groups. Genes common between *Fst* and XPEHH are shown in red.

Most of the common genes on chr5 can be directly related to various stress responses induced by high altitude, e.g. hypoxia (20) and thrombosis (21). For example, *CLP1* is linked to cardiac muscle hypertrophy (22), *YPEL4* has a role in pulmonary diseases (23), *P2RX3* and *ENSGALG00000007381* are involved in blood coagulation (24, 25), and *SLC43A3* plays a possible important role in the repair and growth of the lung tissue under oxidative stress (26). *P2RX3* is also involved in the sensory response to cold and heat, while *PGR2/3* has a role in the immune response (24). Other genes, e.g. *RTN4RL2, UBE2L6, TIMM10*, have been found differentially expressed in cells under hypoxic conditions (27-29) with *RTN4RL2* protecting motor neurons against apoptosis (24), which may be an essential adaptation to high altitude-induced hypoxia. *UTP18* has also been found differentially expressed in human hypoxic cardiomyocytes cell line (29) and is involved in the processing of the pre-18S ribosomal RNA (rRNA). The rRNAs are important components of ribosome - the factory for protein biosynthesis (24). *UTP18* is the only gene that was also detected as a candidate in the GEA in relation to minTemp (Table S6).

Some other notable candidates for adaptation to low temperature and/or high altitude, detected with a strong signal (*ZFst* > 8 or |XPEHH_std| > 4), but from a single approach, include: *SDK1 -* which regulate the dendritic spine development and synaptic connectivity (24), *TRIM3* - involved in nervous system development and critical cellular processes such as proliferation, apoptosis, and transcriptional regulation (30), and *ARFIP2* - with a role in autophagy (24).

In the High group, 25 candidate genes are found, but none were common between *Fst* and XPEHH analyses. The only gene that overlaps a strong SSA signal is *TOGARAM1* (|XPEHH_std|> 4) (Figure 5E), involved in the assembly of non-motile cilia, which are essential for cellular signal transduction (31). Heat-shock induces a rapid loss of these important organelles (32), and *TOGARAM1* may play an important adaptive role in alleviating this effect in high temperature conditions.

Ingenuity Pathway Analysis (IPA) of the 184 candidate genes from the Low group indicates enrichment of processes like lipid metabolism, small molecule biochemistry, and molecular transport (Table S7A), which are expected as hypoxia or cold-temperature stresses affect a cascade of biosynthetic and molecular processes (33, 34). Low group candidates are also associated with many cardiotoxicity terms like bradycardia, cardiac arrhythmia, heart failure, congenital heart anomaly and cardiac enlargement – indicating involvement in hypoxia stress response. Contrarily, the genes from the High group show more enrichment for processes related to organismal growth and development (Table S7A). Low group candidates overlapped with many known QTLs, including those related to skin properties, body temperature, blood parameters, abdominal fat, immune response, and production traits. Meanwhile, High group candidates showed overlap with QTLs for feather properties, disease susceptibility and immune response, and Feed Conversion Ratio (FCR) (Figure 3C, Table S5).

GEA outlier SNPs for the minTemp variable are linked to 339 candidate genes, but only three have relatively large environmental correlations (*r ∼* 0.4), *ULK3 -* a regulator of autophagy in response to stress-induced cellular senescence (cell ageing process) (24), *PITX2 -* a transcription factor which promotes heart cell repair in response to ischemic (low blood supply) injury (35), and *CDH8* - involved in response to cold stress (24) (Table S6).

### Adaptation to extreme rainfall patterns

Three of the six environmental parameters found in the ENM are related to rainfall (PrecSeasonality – variation in precipitation across the year, PrecWQ – precipitation during the wettest quarter, and PrecDQ – precipitation during the driest quarter). Precipitation variables can affect chicken biology in different ways, e.g. insufficient rainfall may limit access to drinking water, while excessive rainfall may facilitate the spread of pathogens and parasites, challenging chicken immunity (36).

Analysis of the precipitation variables provided a strong indication of adaptation to restricted water availability as more SSRs (61% of 427 regions) and stronger signals were observed in populations where water scarcity is likely an issue; for example, in agro-ecologies with either a low rainfall (Low groups for precDQ and precWQ) or large seasonal variation in rainfall (High group for precSeasonality) (Figure 3B, Table S3). Prolonged water deprivation causes dehydration, which can have serious consequences on the overall physiology due to shrinkage of cells, salt-water imbalance in the body, increased osmotic pressure, renal dysfunction, and disruption of the temperature regulatory cues in the brain (37, 38).

Analysis of the precipitation variables detected 500 genes overlapping SSRs, with 14 commonly identified in *Fst* and XPEHH analyses (Table 1). Ten of these genes come from the precDQ Low group indicating their importance for the adaptation to dry environments where access to water may be an issue. The other four genes come from the precSeasonality analysis, of which two are from the High group indicating possible association to water restriction. Closer inspection of the High group populations (Meseret and Gijet) for PrecSeasonality confirms not only very little rainfall during the driest season (average 9.45 mm/m^2^) but also much lower rainfall in the wettest season (average 461 mm/m^2^) compared to that of all 25 populations (average 697 mm/m^2^).

The commonly detected *Fst* and XPEHH candidate genes affect many biological processes as expected from water scarcity stress, and they include a number of lncRNA genes with possible cis-regulatory roles on nearby genes (Table 1). Figure 6G highlights the chr3:71840000-73980000 region from precDQ analysis as it harbours a cluster of several candidate genes (n = 8) and some of the strongest signals. SNPs surrounding *MANEA* show the largest allele frequency difference (dAAF). This gene has been related to behavioural issues like panic disorder in human (39), and it is ubiquitously expressed in many tissues, but most prominently in the urinary bladder and the thyroid (40). Another important candidate from the same region is *EPHA7*, which has also been detected in the GEA analysis in relation to the precSeasonality (*r =* 0.32) variable. This gene is involved in many gene ontology (GO) biological processes (24) including apoptotic process, axon guidance (in response to environmental cues), brain development, ephrin receptor signalling (important in kidney physiology), and nephric duct morphogenesis.

**Figure 6:**
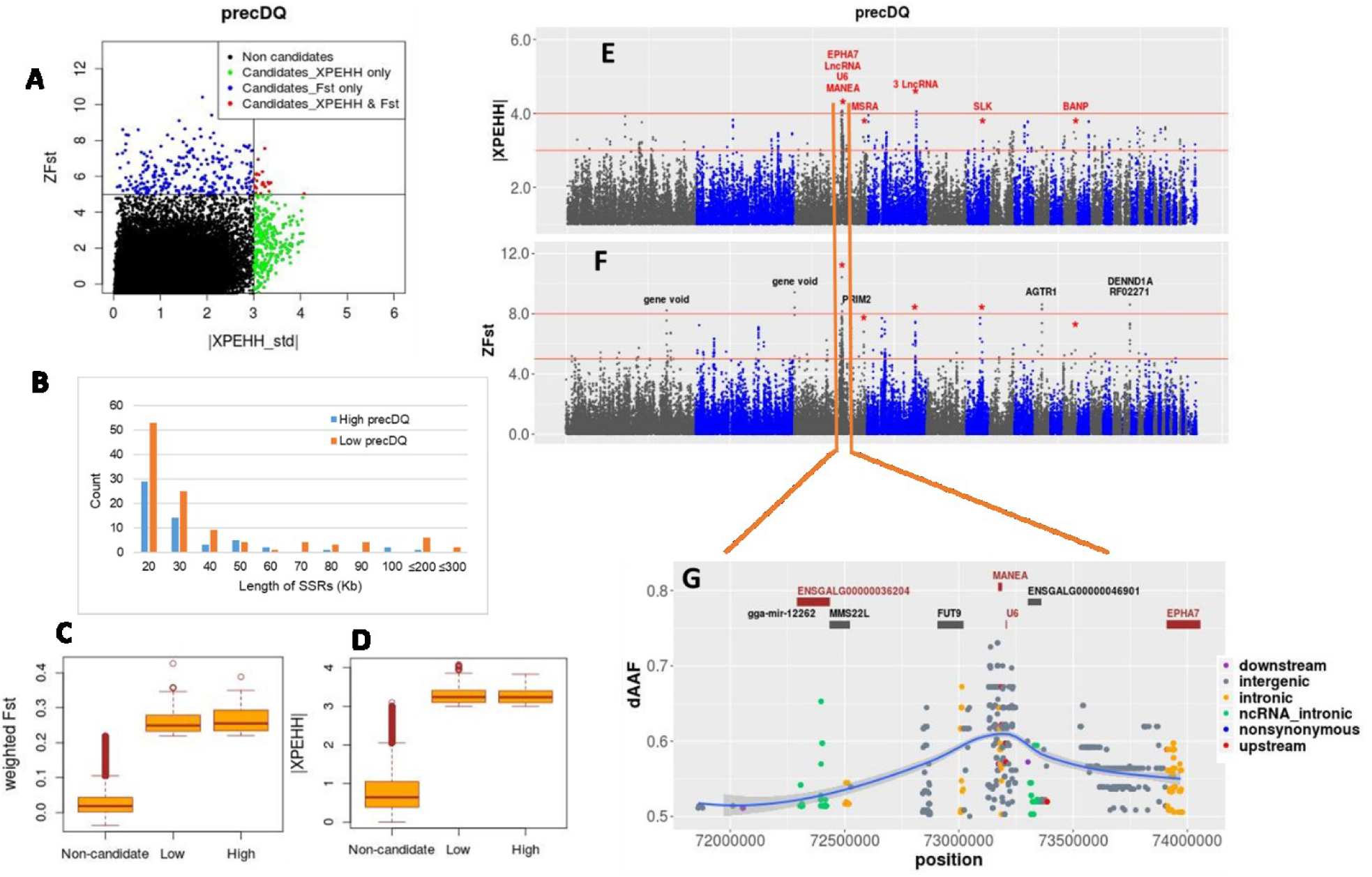
Selection signature analysis results for precDQ. (A) Scatter plot of standardized values of XPEHH versus *Fst*. (B) Length distribution of Selective Sweep Regions (SSRs). (C-D) Box plots showing the distribution of *Fst* and XPEHH metrics for non-candidate and candidate windows. (E-F) Manhattan plots for the XPEHH and *Fst* analyses; common windows are marked with asterisk and gene names from common windows are shown in red. (G) Closer look of the common *Fst*/XPEHH region – chr3: 71840000-73980000 with SNPs showing allele frequency difference (dAAF) > 0.5 between the Low (Gijet, Kido) and High (Kumato, Loya) groups. Genes common between *Fst* and XPEHH are shown in red.

*HTR2C* is another common candidate gene from *Fst* and XPEHH (Table 1) that has also been detected in the GEA. It is associated with the precSeasonality (r= 0.34) environmental variable. This gene plays a crucial mediatory role in the stress-induced activation of the hypothalamic-pituitary-adrenal (HPA) axis (41). It is involved in many biological processes, including behavioural fear response, regulation of appetite, and regulation of nervous-system processes (24).

Genes that overlap with strong signals from a single approach (*ZFst* > 8 or |XPEHH_std| > 4), in relation to scarcity of water, include *AGTR1, TMEM206* and *ATF3. AGTR1* plays a role in the regulation of blood pressure, sodium retention by the kidney, and in kidney development (24). *TMEM206* is involved in pH-gated chloride channel activity that helps to maintain the body’s acid-base balance (24). *ATF3* is a previously reported common stress-responsive transcription factor (24, 42).

We detected two common candidate genes from the precSeasonality Low group, i.e. the population showing low variation in rainfall pattern. *CACNB4* has a functional role in the calcium ion transport (24), whereas *ENSGALG00000053888* encodes a lncRNA with possible regulatory function. It is not clear what type of stress response these genes are involved with, but these may be associated with the adaptation to an environment with high ambient humidity. We observe that the relative humidity in the precSeasonality Low group (59 - 109%) is generally higher both in the wettest and driest quarters of a year compared to that in the High group (57 - 71%) (Figure S11). Two other genes overlap with strong signals in populations experiencing greater rainfall or lower variation in annual rainfall: *PRIM2* (from the precDQ High group; *ZFst* > 8) with a role in DNA replication and *GTDC1* (from the precSeasonality Low group; |XPEHH_std|> 4) showing ubiquitous expression among tissues (24). IPA analyses of the candidate genes from both the Low and High groups (Tables S7B-7D) as well as the overlap of the genes with known QTLs (Figure 3C, Table S5) indicate their involvement in a wide range of biological processes.

The GEA outlier SNPs identified 822 candidate genes in relation to the precipitation variables. Only 33 (4%) of these genes are linked to SNPs with r ≥ 0.4. These genes are involved in a wide variety of physiological and molecular processes such as involvement in stress responses (*PRR5L, ST3GAL6, WIF1, RELL1)*, nervous system processes (*LRRC7, FLOT2*), immune responses (*LCP2, SUSD4*), proteolytic functions (*CPQ, ADAM33*, several Zinc-Finger proteins) which are an important adaptive mechanism to the water deficit condition (43), and genes with potential transcriptional regulatory functions (*ZNF800, SUPT3H*, and several lncRNA genes) (Table S8).

### Soil organic carbon an indicator of source and abundance of food for scavenging chickens

Soil organic carbon (SoilOrgC) affects the nature and abundance of animal biomass in the soil. A High SoilOrgC will be characterized by the presence of many earth-dwelling organisms such as insects and worms that are excellent sources of protein-rich food for chickens. Soils rich in organic carbon also provide fertile ground for wild vegetation and for growing crops - equally important for scavenging birds. Only two chromosomal regions (chr1: 197270000-197290000 and chr10: 6820000-6850000) and seven genes – all in the High SoilOrgC group - were identified in both *Fst* and XPEHH analyses (Figure 7). The SSR in chr1 overlaps with a cluster of several heme-binding genes (*HBBA, HBE, HBE1* and *HBBR*) involved in oxygen carrier activity. The *HBE1* gene is also involved in GO biological processes: response to organic cyclic compound and protein hetero-oligomerisation (24). The dAAF pattern is mostly homogeneous across these genes, but a few intronic SNPs from *HBE1* show the largest dAAF (Figure 7G). These genes are overexpressed in broilers with severe myopathic breast muscles, possibly as a response to insufficient oxygen and oxidative stress (44). Accordingly, their expression may be adaptive in scavenging chickens in response to oxidative stress from food conversion in high SoilOrgC agro-ecology.

**Figure 7:**
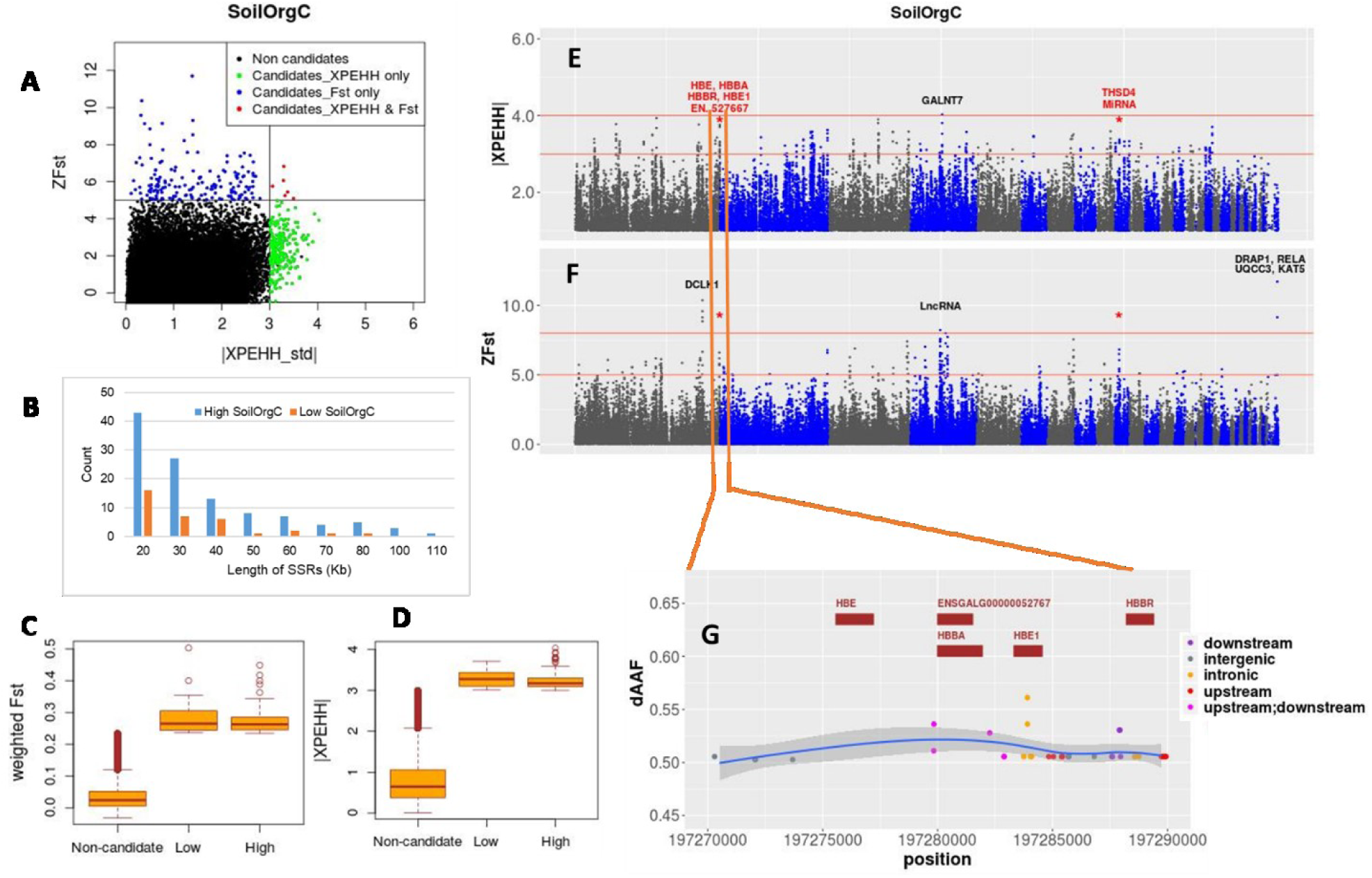
Selection signature analysis results for SoilOrgC. (A) Scatter plot of standardized values of XPEHH versus *Fst*. (B) Length distribution of Selective Sweep Regions (SSRs). (C-D) Box plots showing the distribution of *Fst* and XPEHH metrics for non-candidate and candidate windows. (E-F) Manhattan plots for the XPEHH and *Fst* analyses; common windows are marked with asterisk and gene names from common windows are shown in red. (G) Closer look of the common *Fst*/XPEHH region chr1:197270000-197290000-with SNPs showing allele frequency difference (dAAF) > 0.5 between the Low (Loya, Kumato) and High (Meseret, Gijet) groups. Genes common between *Fst* and XPEHH are shown in red.

The chr10 region overlaps with two genes, *THSD4* and *ENSGALG00000053176 (miRNA). THSD4* has peptidase activity, and the gene has been found in an *Fst*-based selective sweep between Red Junglefowl (RJF) and commercial birds in a previous study (45). *THSD4* is also a candidate gene for a feed efficiency trait in dairy cattle (46). It is likely that the peptidase activity of *THSD4* plays an essential role in the metabolism of protein-rich foods available in high SoilOrgC agro-ecologies. Located within *THSD4*, the miRNA gene may be postulated to have a regulatory effect on its expression.

Other candidates overlapping strong signals from a single approach in the High SoilOrgC group include *DCLK1* (*ZFst* > 10) with protein kinase activity and possible involvement in the nervous system and forebrain development and *GALNT7* (|XPEHH_std|> 4), involved in carbohydrate metabolic process (24). From the Low SoilOrgC group, four genes overlapped with a strong SSR on chr33 (chr33: 6470000-6500000) (*ZFst*: 9 - 12). These include *DRAP1 -* involved in transcriptional regulation (24), *RELA -* a ubiquitously present transcription factor affecting many biological processes including cell growth, immunity, and apoptosis (24), *UQCC3* which plays an important role in ATP production by mitochondria (24) and *KAT5 -* regulating many biological processes including autophagy under starvation condition (24).

The major candidates from GEA (*r* ≥ 0.4) show a predominant involvement in the brain and the neuronal development and processes (e.g. *KIF5C, ZEB2* (also detected in XPEHH in Low group), *ZNF536, CDH2, FOXO3, FARP, ADGRL3*), which may be linked to the scavenging behaviour of chicken when they actively search and catch food. Another candidate, *C8orf37*, is involved in the morphogenesis of photoreceptor cells of the eye (important for catching prey) and two candidate genes - *RAD54B* and *URI1 -* are related to growth (Table S8).

### Land use pattern is an important determinant of food abundance for scavenging chickens

By-products from the harvesting and processing of cultivated grains or crops are important sources of plant-based food for scavenging chickens (47). Interestingly, from 191 genes overlapping candidate SSRs, the majority (72%) are selected in the Low LandUse group (i.e. populations living in regions with a low proportion of cultivated land), where supplementation with crops residues and grain is expected to be less. Only seven genes from five sweep regions are common to *Fst* and XPEHH analyses, of which six are from the Low group (Table 1). The Low group genes include two clusters: *STS and PUDP* from chr1:127800000-127830000, and *SMYD3* and *KIF26B* from chr3:34260000-34280000. *STS*, with its role in lipid and steroid metabolism, is a candidate gene for growth and feed efficiency traits in cattle (48), while *PUDP* is involved in nucleotide metabolism (24). In the second cluster, *SMYD3* is involved in transcriptional regulation and cellular response to dexamethasone (a corticosteroid hormone) that can affect appetite (49), and *KIF26B* has a role in cell-signalling (24). Another common gene from the Low group is *NDST4*, which has a strong association with low abdominal fat content in chicken (50). *NCAPG* is the only common gene between the *Fst* and XPEHH analyses in the High group (i.e. populations living in regions with a high proportion of cultivated land). This gene has been reported to be associated with various growth-related traits in beef cattle and carcass traits in chicken (51, 52).

From the Low LandUse group, several genes overlapped with strong signalling SSRs from a single approach (*ZFst* > 8 or |XPEHH_std|> 4), including *AGMO* which has roles in lipid metabolism and feed efficiency in chicken (53), *MED8* – involved in transcriptional regulation, and *SZT2* – involved in cellular response to amino-acid and glucose starvation (24). From the High LandUse group, notable genes overlapping with strong SSR signals include *ADIPOR2* – regulating glucose and lipid metabolism, *SNX10*with roles in gastric acid secretion, bone resorption and calcium ion homeostasis, and several genes involved in transcriptional regulation (*PRDM5, CBX3*, and *HNRNPA2B1*) (24). Some of the largest SSRs (300 Kb -550 Kb) were detected in LandUse analysis. These large SSRs predominantly overlap with ncRNA genes with possible regulatory roles.

Strong candidates from the GEA (*r* ≥ 0.4) analysis include a Tyrosine Hydroxylase (*TH*) like gene - with possible roles in regulating the eating behaviour in response to nutrient level, *WASHC3* with a role in protein transportation (also detected as candidate by *Fst* in High group), and two lncRNA genes with possible cis-regulatory effect on nearby genes (Table S8).

### Shared candidates between environmental predictors

We detected between 150 and 219 genes overlapping sweep regions in relation to each environmental predictor (Figure 3A). Interestingly, about 15% of these genes (152 of 1008) are common to two or more environmental analyses (Figure S17). The proportion of shared genes between environmental analyses varied between 0.3% and 14%. While these shared genes may represent pleiotropic effects or shared pathways of stress response as observed with IPA analysis (Figure S17C), the low rate of overlap in general indicates that our environmental analyses have captured different components of environmental selective pressures that in turn have shaped genomes distinctively.

In a few cases, the same gene was candidate in both Low and High groups for the same environmental analysis, e.g. *ZNF451* (precDQ), and four genes (*ADGRL3, CIS, CIR, HMGCLL1*) for the SoilOrgC parameter. These genes traverse multiple windows, with separate windows under selection in the two groups. Selection of different regulatory elements or use of alternatively spliced transcripts of the genes in the opposing groups may be responsible for the results. However, while intronic variants from these genes show generally the largest dAAFs, none are annotated as splice variants.

## Discussion

Local environmental adaptation, demonstrated by indigenous livestock populations, is an important resource for promoting sustainable breeding improvement in response to the challenges arising from changes in climatic conditions, production environments, and consumer demands. Knowledge regarding the genetic basis of such adaptation will allow targeting of favourable genes in breeding programmes alongside the productivity traits using one or more genetic interventions (e.g. marker-assisted selection, genomic selection and/or gene editing). Dissecting the genetics underlying environmental adaptation, however, is difficult due to the complexity of agro-climatic stressors posing as selection pressure.

To address this issue, our study applied a powerful interdisciplinary approach to first, disentangle and identify the important agro-climatic drivers of adaptation from a large array of environmental parameters and then to identify the associated candidate regions, their genes, and variants using two complementary approaches (SSA and GEA). In the *Fst* and XPEHH-based selection signature analyses, we compared extreme populations in relation to single environmental parameters, while in the RDA analysis we looked for genotype-environment association across all samples and all environmental predictors simultaneously. This is the first study to the best of our knowledge to perform such rigorous and comprehensive analysis on a livestock species and more specifically on chicken.

Our study has detected strong adaptive signals in relation to altitude and climate-driven stresses (hypoxia, thrombosis, low temperature, and water scarcity) and, for the first time, identified the environmental proxies associated with the genetic adaptations linked to scavenging conditions, reflecting the impact of the nature and abundance of food on indigenous chickens. While we detected strong adaptive signals linked to survivability at high altitude and in cold temperature, we found few strong selective signals in relation to high-temperature stress in Ethiopian chickens. It is a possible legacy of the origin and history of African indigenous chickens. Indeed, the Red Junglefowl, the main ancestor of the domestic chicken (54), with native habitat in tropical South and South-East, is adapted to the hot and humid climates of these regions (55). Accordingly, it may be expected that the chickens introduced to Africa were already adapted to the high temperature witnessed by the Ethiopian populations studied.

Comparison of our results with the few other available adaptation studies on chicken found little overlap among the candidate genes. For instance, similar to our study, Zhang *et al*. (9) reported candidate genes affecting cardiovascular and respiratory systems, and immune responses in Tibetan highland chickens as adaptations to hypoxia, but none are common with our set. This result may be attributed to a number of factors. First, rapid adaptation often works on the “standing variations” in a population (56). Different demographic history of the African and Tibetan chickens may have offered distinct standing variations for the natural selection to work on. As a result, even though the adaptive responses were similar in both instances, the associated genes were different. Moreover, epistatic interactions and pleiotropic effects of genes may favour selection of one gene over another in different geographic areas (57). Selection of different genes in different populations in relation to high altitude has also been observed in human studies (56). These results illustrate the plasticity of the genome in its response to environmental selection pressures.

Among the identified candidate genes associated with each environmental predictor, only a few may be considered as strong candidates being detected by multiple approaches or coinciding with extreme signals from a single approach. It supports that environmental adaptations observed in Ethiopian indigenous chickens are predominantly under oligogenic control, as supported by the results of other studies (56, 58). Also in many instances, we find important biologically relevant candidate genes residing in the same selection region or at close proximity (e.g. we detected nine major candidates associated with hypoxia, low temperature, and thrombosis from a single SSR in chr5, and four heme-binding genes detected as a cluster from the SoilOrgC analysis in chr1). Such clusters of genes may actually be at the root of rapid adaptation to extreme environment as these clusters may be under the genetic control of one or a few regulatory variants only, thereby allowing the natural selection to achieve rapid genomic response in relation to new environmental challenges. While further deeper investigation are required, including the identification of the causative mutations, such results provide a new framework to explain the rapid adaptation and success of an ubiquitously adapted species like chicken to different agro-ecologies.

Our findings have also direct implication for achieving fast and sustainable improvements in new breeding programmes aiming to produce chicken lines that will be both productive and well adapted to the African backyard farming system. Given the oligogenic regulation of adaptive traits, the best option for achieving genetic progress through breeding improvement program would be to combine Genomic Estimated Breeding Value (GEBV) for production traits with a targeted marker-associated selection for the environmentally adaptive regions of the genome.

Finally, our study exemplifies the use of ENM as a powerful predictive tool for adaptation studies in livestock. Uniquely, it allowed us to first identify the drivers of environmental selection and then to investigate the corresponding genome responses, instead of the contrary approach commonly adopted in animal adaptation studies. Moreover, we envisage many other different applications of ENM for livestock. For example, environmental characterization of populations, as shown in the present study, can be the basis for characterizing or even defining livestock ecotypes. Along with the agro-climatic data, any other environmental variable (e.g. epidemiological) may be included (59). Habitat suitability mapping can be a useful approach for predicting suitable areas for introducing exotic breeds (60) or for predicting the impact of climatic change on livestock habitats. These can be extremely valuable in conservation of important livestock genetic resources to meet future demand.

## Methods

### Sampling design

Chicken samples analyzed here consisted of 225 birds from 23 populations (villages or Kebeles), collected for this study and 20 samples from two other populations (Horro and Jarso) from a previous study (54). The sampled populations represent different agro-ecological zones (61) distributed across 13 districts of Ethiopia and across six of the nine national regional states (Afar, Amhara, Gumuz, Oromia, SNNPR (Southern Nations, Nationalities, and People’s Region), and Tigray) (Table S1). Blood samples from 8-10 chickens per population were collected for genomic analysis. Sampling was performed with the logistical support and agreement of the Ethiopian Ministry of Agriculture and Ethiopian Institute of Agricultural Research (EIAR). All animal works were approved by the Institutional Animal Care and Use Committee of the International Livestock Research Institute (IREC2017-26). Geographic coordinates (latitude and longitude) of the sampling villages were recorded, providing the entry points for the collection of environmental data for ENM.

### Environmental data

Environmental data across the Ethiopian landscape were obtained from several public databases at a spatial resolution of 30 arc-seconds (∼1 km^2^) (Table S2). Climatic data were extracted from the WorldClim database (v1 and v2) (62). Soil property data were downloaded from the SoilGrids1km v0.5.8 database (63) and the Spatial Data Access Tool from NASA (64). Data on land use and land cover were obtained from the Harmonized World Soil Database V1.2 (65). The Crop_dominance variable was accessed from the Global Food Security Analysis-Support Data (66).

For each population, we originally recorded a single central coordinate (e.g. market place) in the village. However, to capture the environmental condition for the village, we selected nine additional geographic coordinates in separate grids surrounding the actual recorded location for each population. These grids were within 1.2 km from each other and were located using Google Earth Pro v7.3. The environmental data were extracted using the ‘raster’ R package, resulting in a total of 250 coordinate points from all 25 populations and these were considered as “occurrence points” for ENM.

### Ecological Niche Modelling

ENM was performed using MaxEnt v3.4.1 (13). To avoid overfitting of the models, the environmental parameters were first shortlisted using the R package ‘MaxentVariableSelection’ (67) which helped remove correlated variables and selected those with large contributions. The best combination of model parameters was chosen by running MaxEnt with different combinations of Feature Classes (FCs: Linear, Quadratic, Product, Hinge, Categorical, and Threshold) and Regularisation Multiplier values (RM: 0.1 to 6 with steps of 0.5), and by evaluating the performance of these models based on AICc (**A**kaike **I**nformation **C**riterion **c**orrected for small samples) values with ENMeval (68). The final models were run using the Hinge, Quadratic and Product FCs and RM of 3.5 as this combination was found to be the best with the lowest AICc value (Figure S21). Random*k*fold (with k = 10) method was used to partition “occurrence” data for training and testing purposes in ENMeval. The predictive power of the models was assessed using the Area Under ROC Curve (AUC) values where AUC > 0.5 was considered as higher predictive power, AUC = 0.5 as random predictors, while values < 0.5 as worse than random predictor (69) (Figure S22A). A jackknife assay was performed for the test and training data to assess the importance of the variables (Figure S22B). Habitat suitability maps were generated using MaxEnt’s cumulative output.

### Whole-genome sequencing and data processing

WGS was performed on an Illumina HiSeqX platform in paired-end mode with a read length of 150 bp and average coverage of ∼40X. Sequence reads were mapped against the GRCg6a reference assembly using BWA-mem (v. 0.7.15-r1140 (70) Variant calling was performed following the GATK v3.4 best practice protocol (71) involving the Haplotype Caller method and Joint Genotyping of all samples together. Initial variant filtration was performed using the VQSR approach (72) in GTAK using 1 M validated SNPs (73) and over 20 M known chicken SNPs from Ensembl release 92 (the variant coordinates were converted from Gal5 to GRCg6a by mapping with flanking sequence).

For downstream genomic analyses, only autosomal variants were used, and any individuals with high relatedness (co-efficient > 0.9) were removed. SNPs that did not pass the following criteria were excluded: genotype quality ≥ 15, depth of coverage ≥ 3 and missing genotype rate < 20%. Relatedness between samples was calculated in VCFtools v0.1.15 (74) using the --relatedness option. Nucleotide diversity was calculated using the --site-pi option in VCFtools.

PCA was performed using the Eigenstrat method in Eigensoft v6.1.4 software (75). Admixture analysis was performed in ADMIXTURE programme v1.3.0 (76) with *K* values 1-5, and the best *K* value was chosen based on the cross-validation method.

### Selection signature analysis

Extreme Low and High groups of populations were chosen by ranking the 25 populations for each environmental parameter and selecting two populations from each end of the gradations. SSA was performed in overlapping sliding windows with at least 10 SNPs. *Fst* analyses were performed in VCFtools using the Weir and Cockerham approach (14). The weighted *Fst* values were standardized (*ZFst*) to allow setting the same threshold across analyses.

XPEHH analyses were carried out using the Hapbin package (77) after removing any SNPs with missing genotypes. The genetic map positions for SNPs were calculated using chromosome-specific recombination rates based on Groenen *et al*. (78) except for chr16 for which recombination rate was estimated based on Elferink *et al*. (79). Genotype data were phased in Beagle v5.1 (80). XPEHH analyses were first performed for individual SNPs by setting options for minor allele frequency to be 5% and the cut-off value for Extended Haplotype Homozygosity (EHH) to be 0.1. Subsequently, mean values were calculated within windows for both the standardized XPEHH (XPEHH_std) and the absolute value of XPEHH_std. Only windows with all the SNPs showing the same directionality of selection were considered for sweep detection, with positive XPEHH values indicating selection in the Low group, and negative values representing selection in the High group. Fewer windows were analyzed in XPEHH (91.2K - 91.6K) compared to *Fst* (92.4K - 92.5K), following the application of the extra filtrations steps. Empirical *P*-values were calculated for both *Fst* and XPEHH by ranking the windows based on each metric.

Moreover, pooled heterozygosity (*Hp*) (16) in windows was calculated for Low and High groups separately to provide an extra source of support for the directionality of selection. *Hp* results were consulted when directionality could not be established from the XPEHH result unambiguously or because those windows were not analyzed in XPEHH. Windows were removed from the downstream analysis when the directionality could not be resolved.

### Genotype-environment association analyses

RDA was performed in Vegan v2.5-4 in R (81) following Forester (2019) (82). Autosomal SNPs (without any missing genotypes) were first LD pruned using PLINK v1.9 with the “--indep-pairwise” command with window size = 10kb, step size = 10 SNPs and r^2^ > 0.5. The genotype data of the remaining sites were converted as dosage (number) of non-reference allele (0, 1 and 2). RDA was run with genotypes as the response variables and environmental data as the explanatory variables conditioned on latitude and longitude.

Significance of the overall model and the RDA axes were estimated by the anova.cca function in Vegan with permutation numbers of 499 and 49, respectively. Given the normal distribution of SNP loadings, a cut-off value of SD >= 3.5 was used for detecting outliers (two-tailed *P*-value = 0.0005) for each significant axis. SNPs in high LD (r^2^ > 0.8) with outliers were identified by calculating LD in PLINK by specifying the following options: -*-r2 --ld-snp-list <candidate_SNP_list> --ld-window-r2 0*.*8 --ld-window-kb 40kbs*.

### Functional interpretation of SSRs, candidate genes and variants

The putative SSRs were intersected with known genes from Ensembl (release 98) using Bedtools v2.26 (83). Candidate genes were checked for their overlap with known chicken QTLs (18). Only significant QTLs with size < 1 Mb were considered. Candidate genes were also analyzed for their molecular and cellular functions, and physiological processes using IPA (84). SNPs within SSRs were annotated using the programme ANNOVAR (85). Hypergeometric tests for under-or over-representation of the SNP annotation categories were performed in R using the “phyper” function.

## Supporting information

Main Supplementary File

## Acknowledgments

This study was funded by the Bill and Melinda Gates Foundation (BMGF) and with UK aid from the UK Government’s Department for International Development (Grant Agreement OPP1127286) under the auspices of the Centre for Tropical Livestock Genetics and Health (CTLGH), established jointly by the University of Edinburgh, SRUC (Scotland’s Rural College) and the International Livestock Research Institute. The findings and conclusions contained within are those of the authors and do not necessarily reflect positions or policies of neither the BMGF nor the UK Government. This research was conducted as part of the CGIAR Research Program on Livestock and is supported by contributors to the CGIAR Trust Fund. CGIAR is a global research partnership for a food-secure future. Its science is carried out by 15 Research Centres in close collaboration with hundreds of partners across the globe (http://www.cgiar.org/about-us/our-funders/). Adriana Vallejo is a PhD student funded by Vice-Chancellor Scholarship for Research Excellence International at University of Nottingham and Administrative Department of Science, Technology and Innovation (Colciencias) – Colombian Government (Call 2015 N°728). We also thank Prof. Nick Sparks (CTLGH, SRUC) for his valuable support in conducting this research. We would also like to thank Edinburgh Genomics (Edinburgh, UK) for producing the sequence data used in this study.

## Availability of data

Whole genome sequence data have been deposited in the European Nucleotide Archive (ENA) under study accession numbers PRJEB39275.

## Competing interest statement

The authors have no competing interests

## Notes

### Competing Interest Statement

The authors have declared no competing interest.

