## Supplementary material for "Integrated environmental and genomic analysis reveals the drivers of local adaptation in African indigenous chickens": Main Supplementary File

**This PDF file includes:**

Figures S1 to S17

Tables S1 to S8 (Tables S3-S6 are submitted as separate supplementary excel files)

SI References

Supplementary Figures

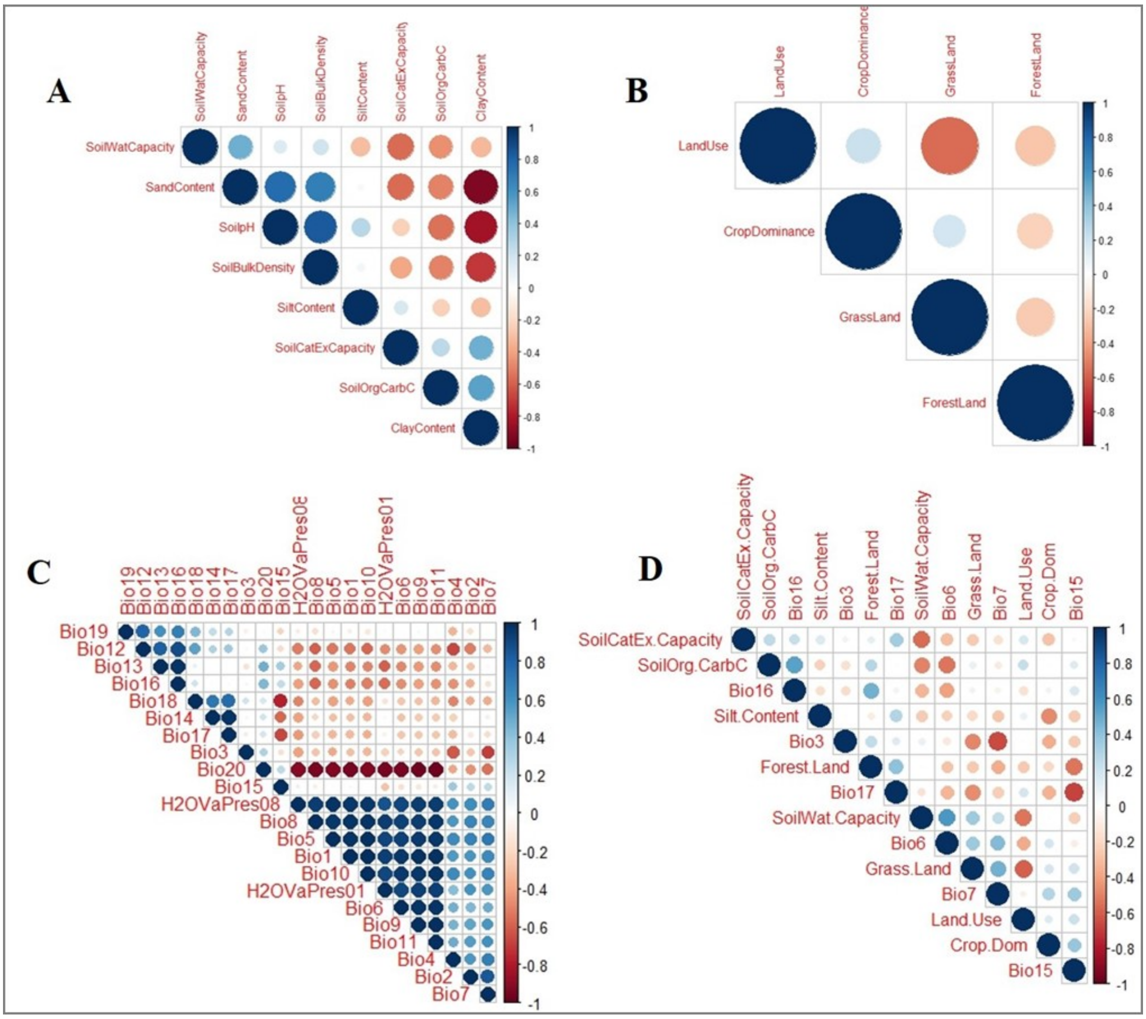


**Figure S1.** Spearman correlation analysis of environmental parameters: (A) Soil variables, (B) Vegetation and land cover variables, (C) Climatic variables, (D) correlation analysis between non-correlated variables from different groups.

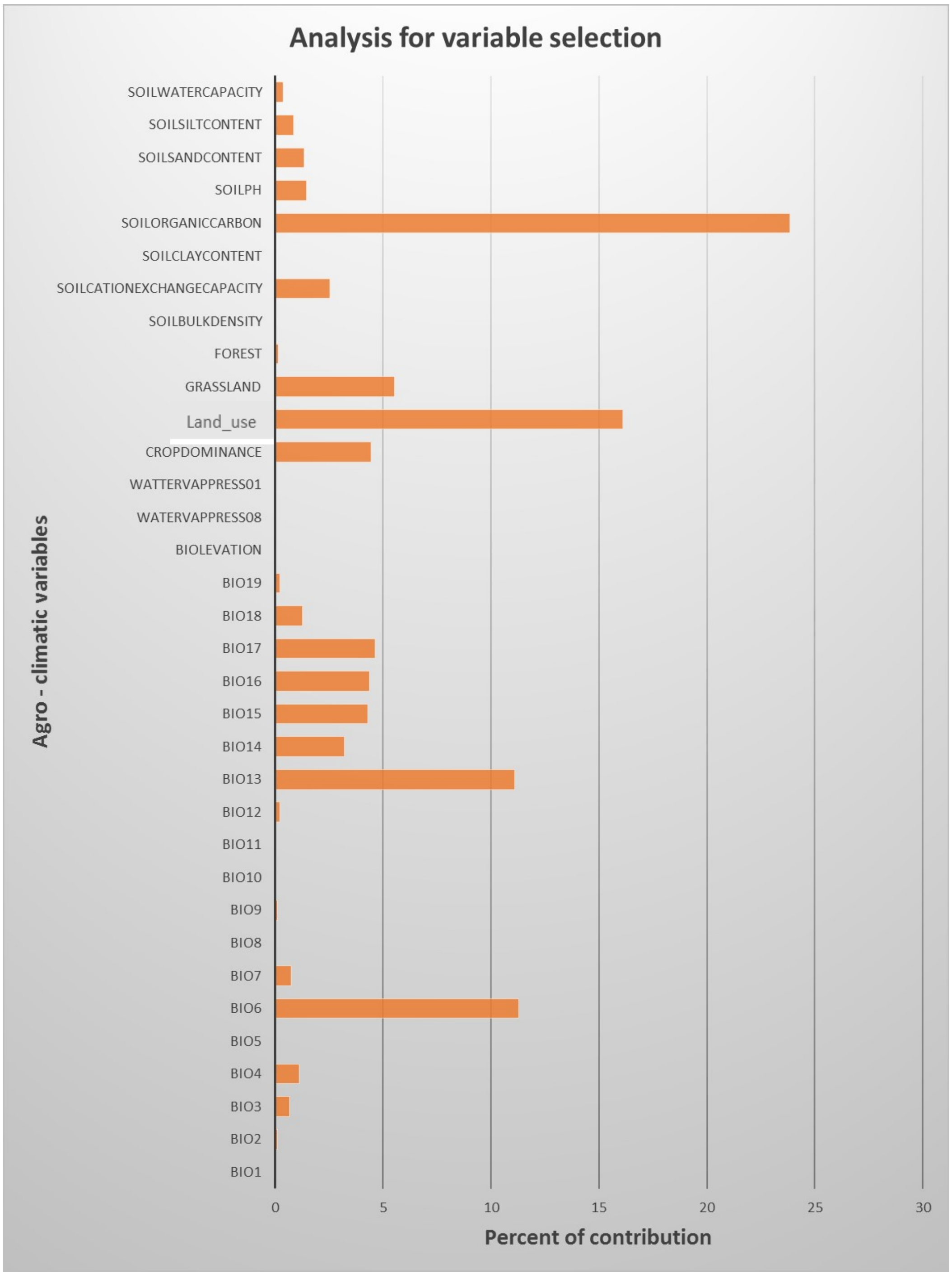


**Figure S2.** Percent contributions of each of the 34 agro-climatic variables calculated using the MaxEntVariableSelection R package. Variables with values >= 4% were retained for the final model

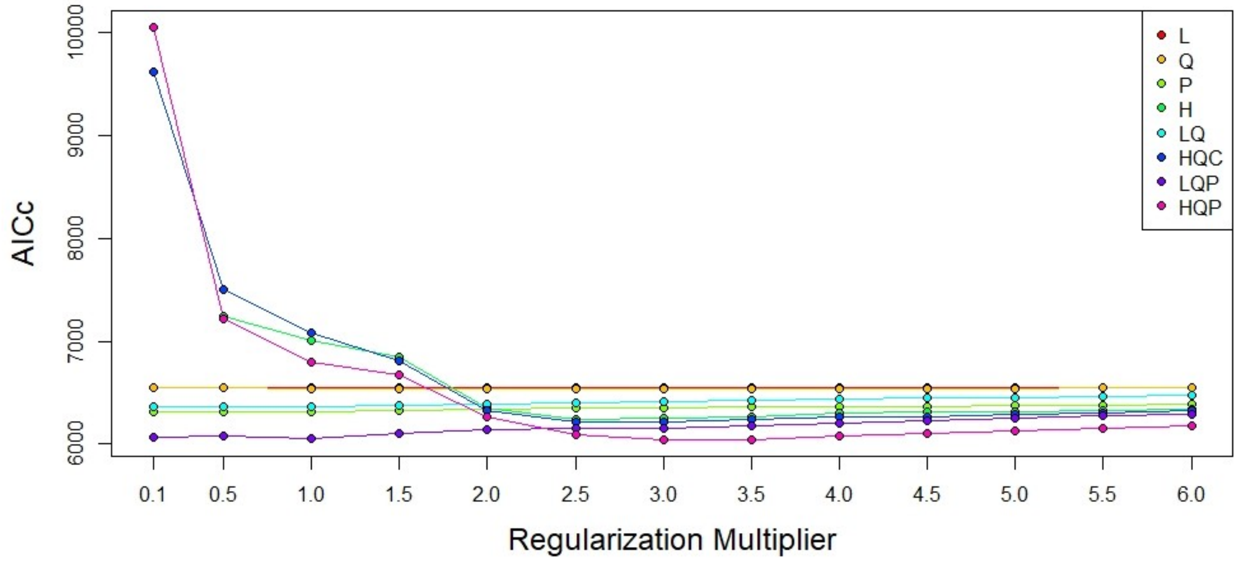


**Figure S3:** AICc values for different combinations of Feature Classes (FCs) and Regularization Multipliers. The FCs shown are: Linear (L), Quadratic (Q), Product (P), Hinge (H) and Categorical (C). The Threshold FC is not depicted here, as it did not show any effect in any combination.

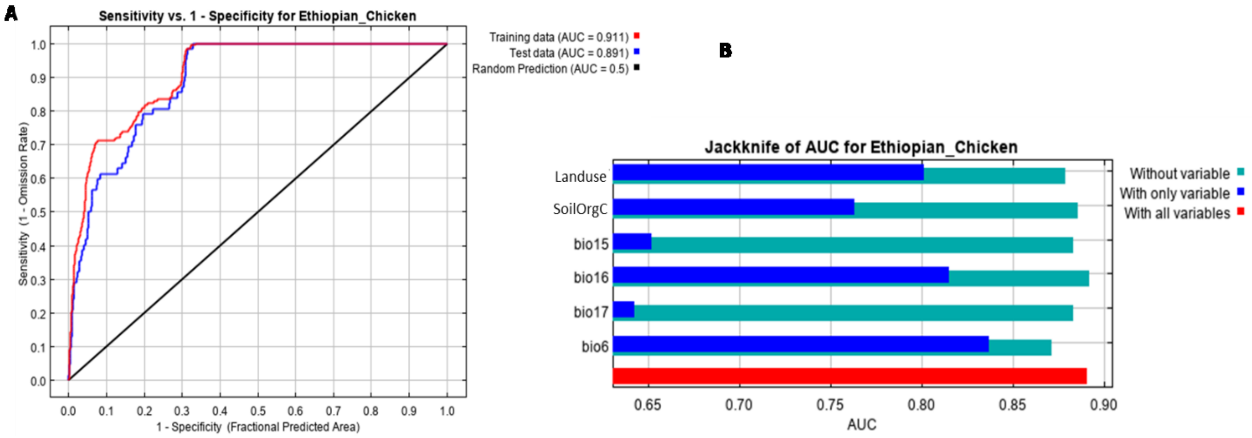


**Figure S4.** MaxEnt model based on the six selected variables (A) Receiver Operating Curve for training and test data, (B) Jackknife result for AUC (Area Under Receiver Operating Curve) for each selected environmental predictors.

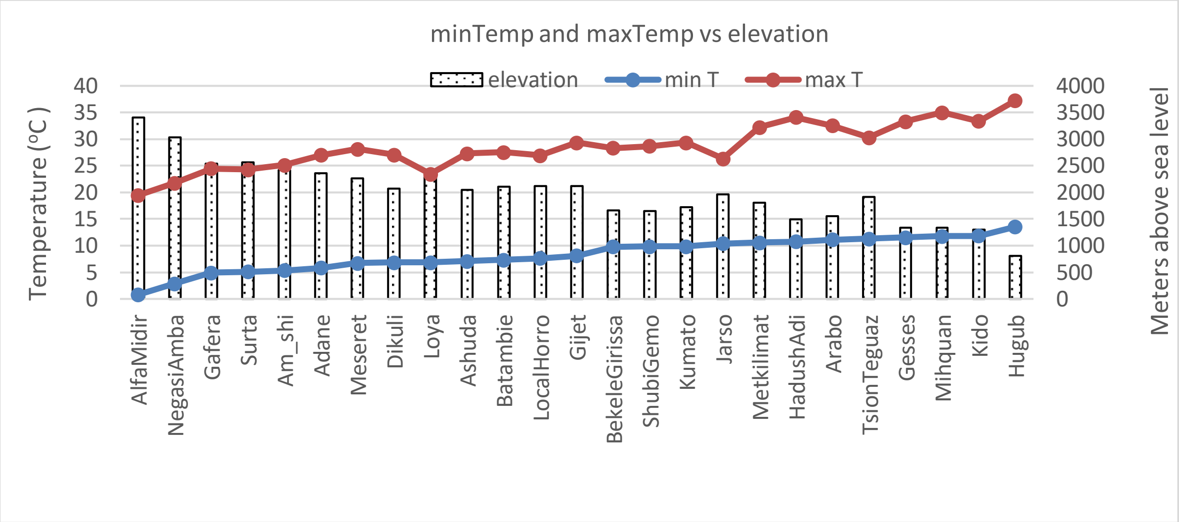


**Figure S5:** Population gradient based on minimum temperature in the coldest month (minTemp). The maximum temperature in the warmest month (maxTemp) and elevation values are also included to show their high positive and negative correlations, respectively with the minTemp parameter. Two populations from each end of the distribution were chosen for selection signature analysis (AlfaMidir and NegasiAmba as Low group and Mihquan and Hugub as High group). For the High group, Mihquan, instead of Kido was chosen as in the former temperatures in both the coldest and warmest months were among the highest and elevation among the lowest compared to other populations.

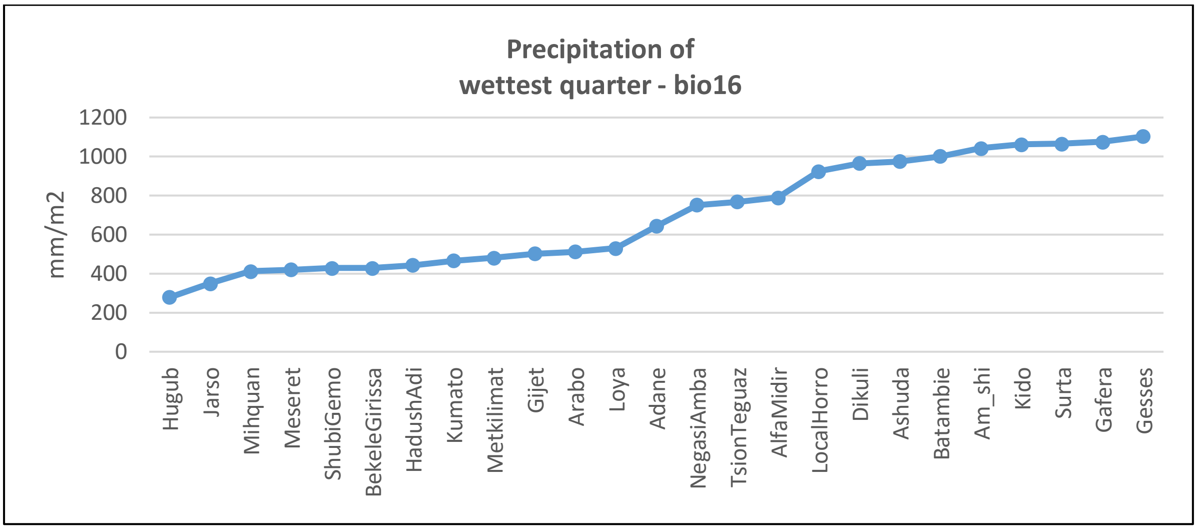


**Figure S6:** Population gradient from low to high based on precipitation of the wettest quarter (precWQ). Two populations from each end of the distribution were chosen for selection signature analysis (Hugub and Jarso as Low group and Gafera and Gesses as High group).

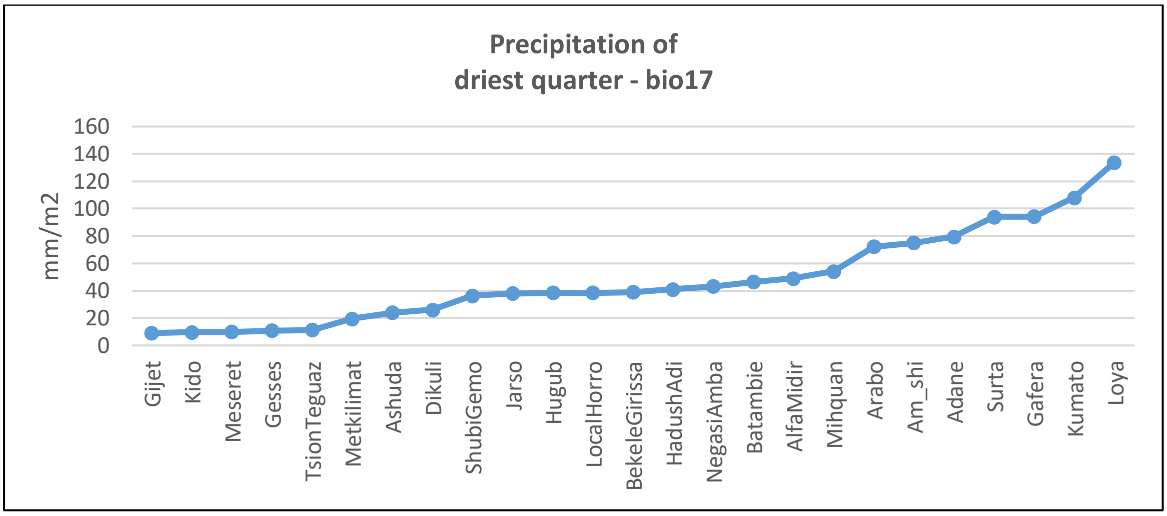


**Figure S7:** Population gradient from low to high based on precipitation of the driest quarter (precDQ). Two populations from each end of the distribution were chosen for selection signature analysis (Gijet and Kido as Low group and Kumato and Loya as High group).

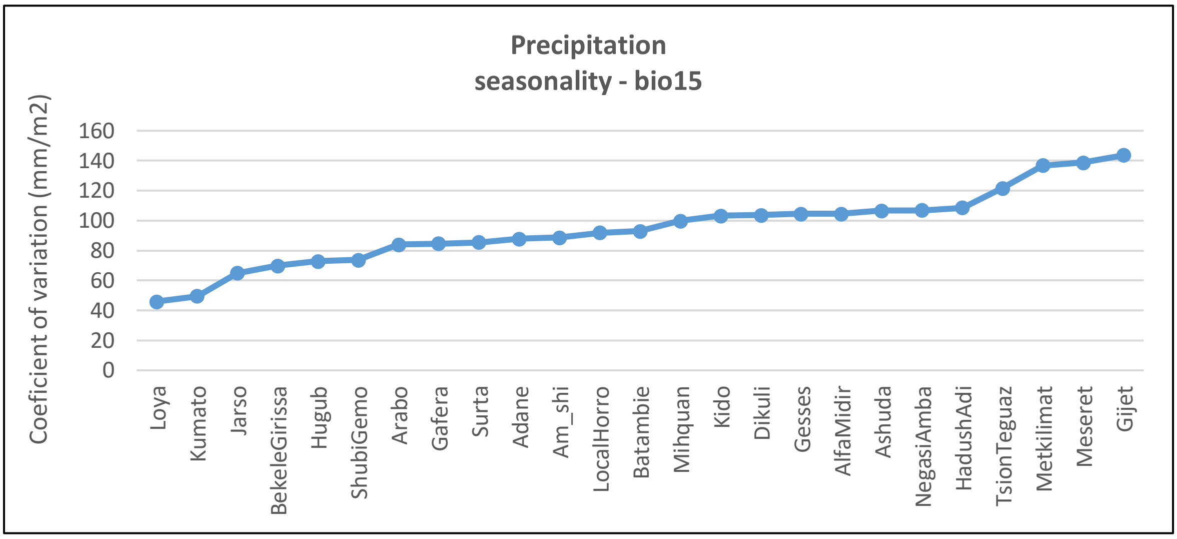


**Figure S8:** Population gradient from low to high based on precipitation seasonality (precSeasonality). Two populations from each end of the distribution were chosen for selection signature analysis (Loya and Kumato as Low group and Meseret and Gijet as High group).

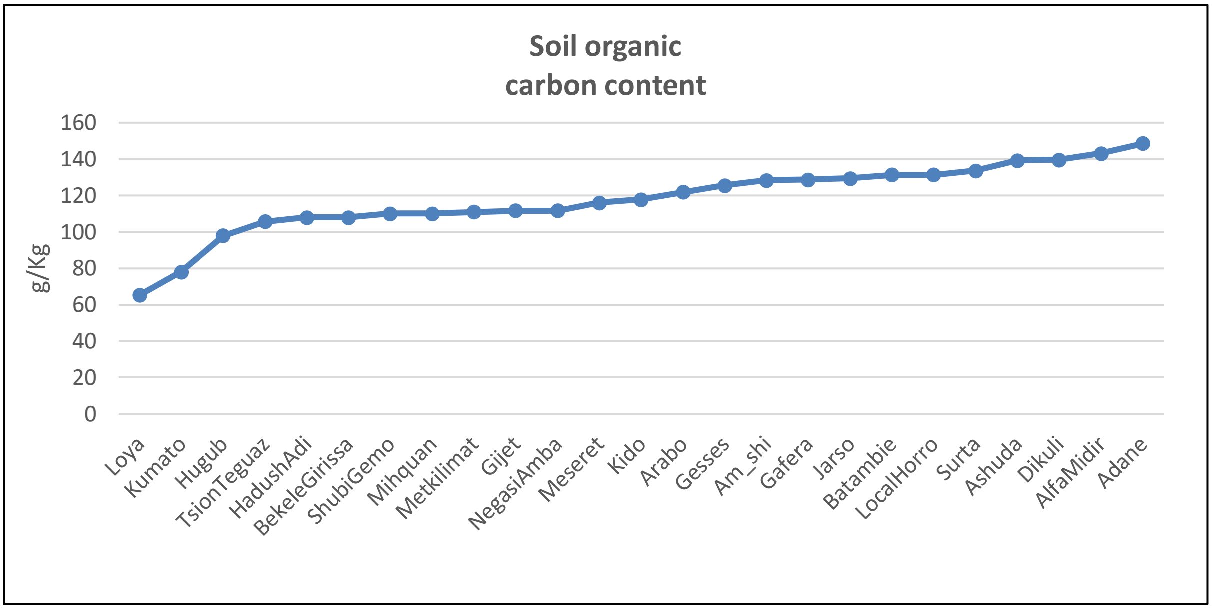


**Figure S9**: Population gradient from low to high based on soil organic carbon content (SoilOrgC). Two populations from each end of the distribution were chosen for selection signature analysis (Loya and Kumato as Low group and AlfaMidir and Adane as High group).

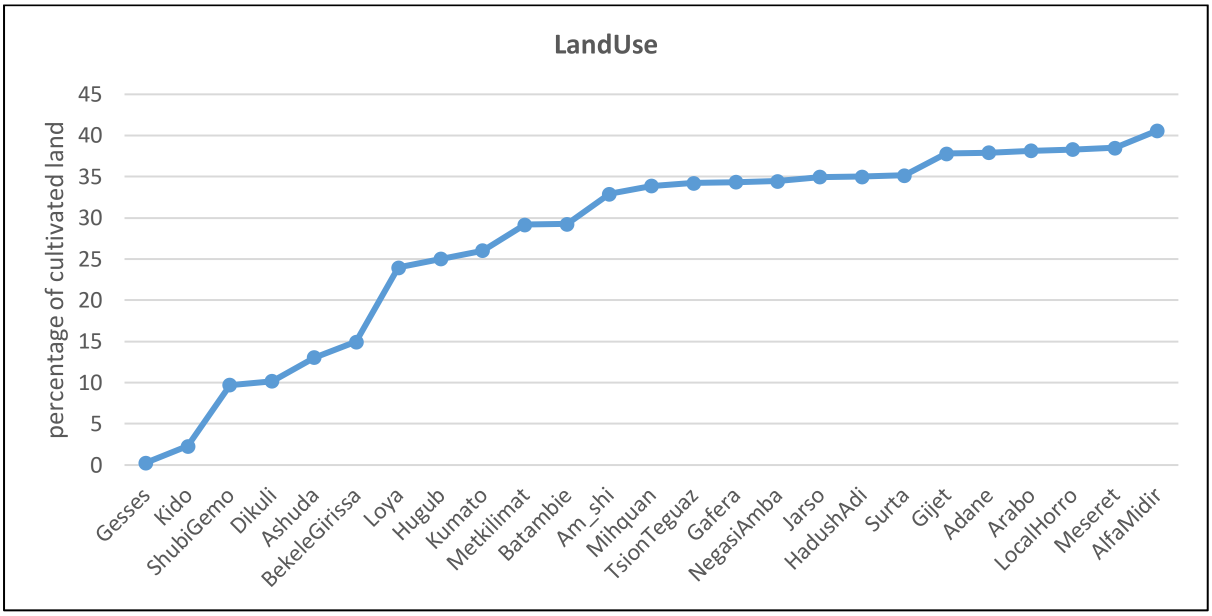


**Figure S10:** Population gradient from low to high based on proportion of cultivated land (LandUse). Two populations from each end of the distribution were chosen for selection signature analysis (Gesses and Kido as Low group and Meseret and AlfaMidir as High group).

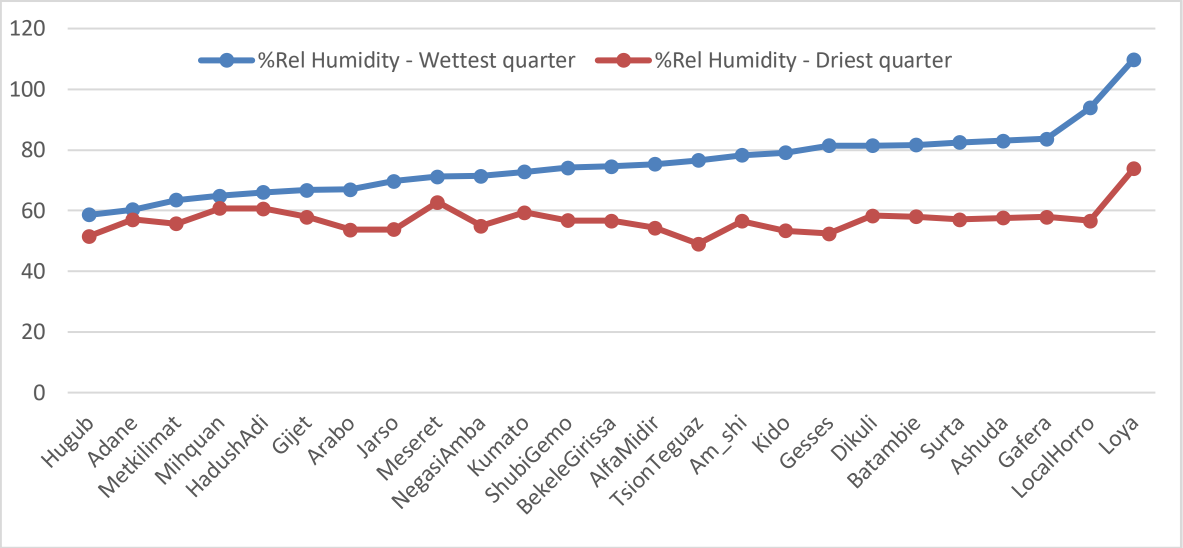


**Figure S11:** Mean Percent relative humidity in different populations in the wettest and driest quarter

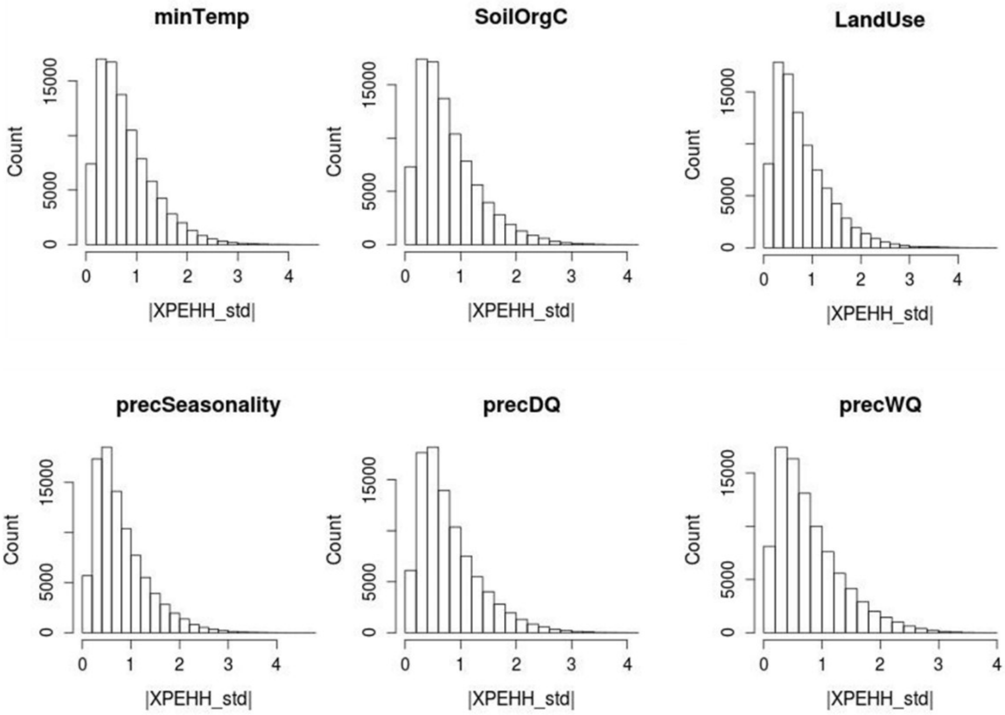


**Figure S12: Histograms of |XPEHH| values in windows in different analyses**

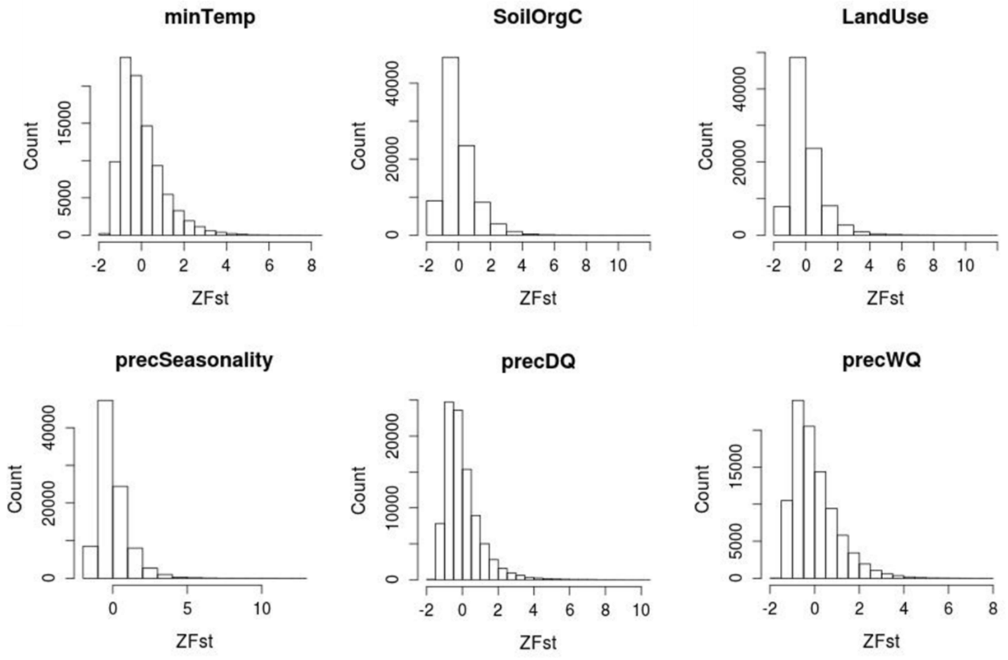


**Figure S13: Histograms of *ZFst* values in windows in different analyses**

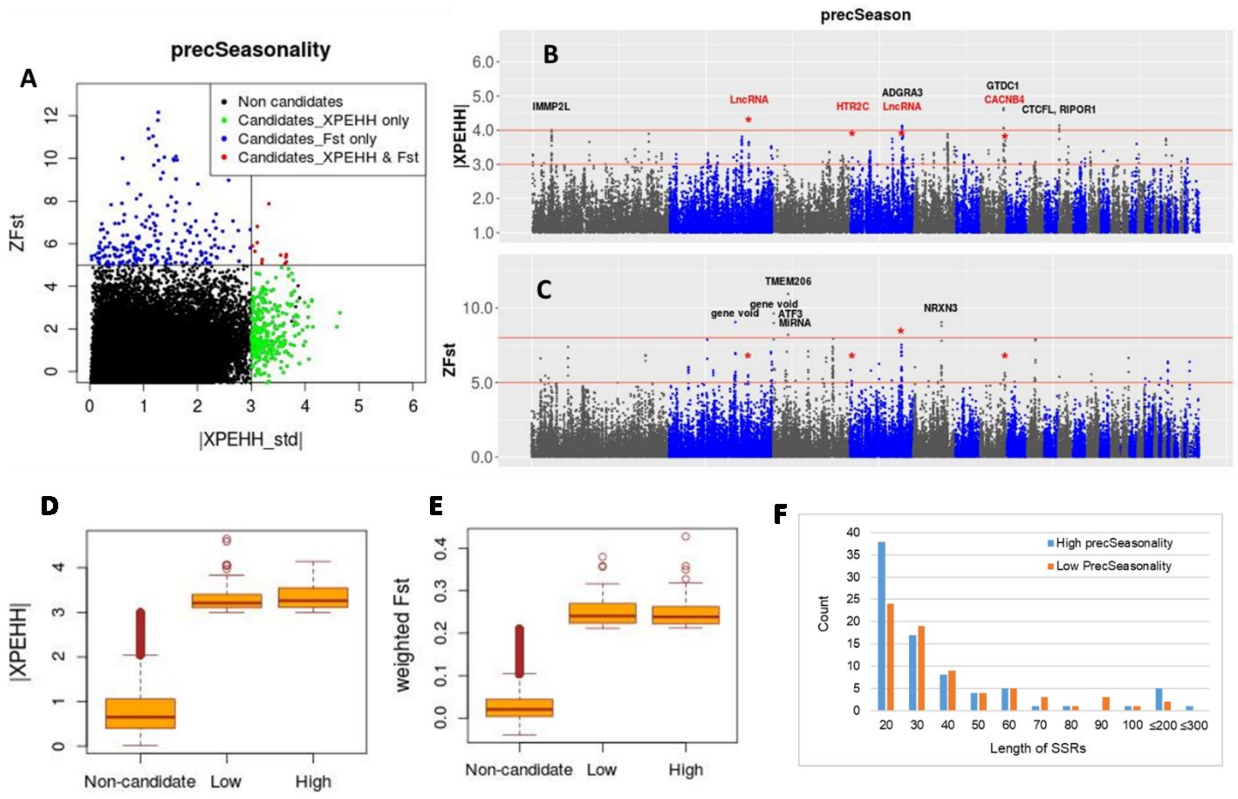


**Figure S14:** Selection signature analysis  results for precSeasonality.  (A) Scatter plot of standardaized values of XPEHH vs *Fst*. (B-C) Manhattan plots for the XPEHH and *Fst* analyses; common windows are marked with asterisk and gene names from common windows are shown in red. (D-E) Box plots showing the distribution of *Fst* and XPEHH metrics for non-candidate and candidate windows from Low and High groups. (F) Length distribution of Selective Sweep Regions (SSRs).

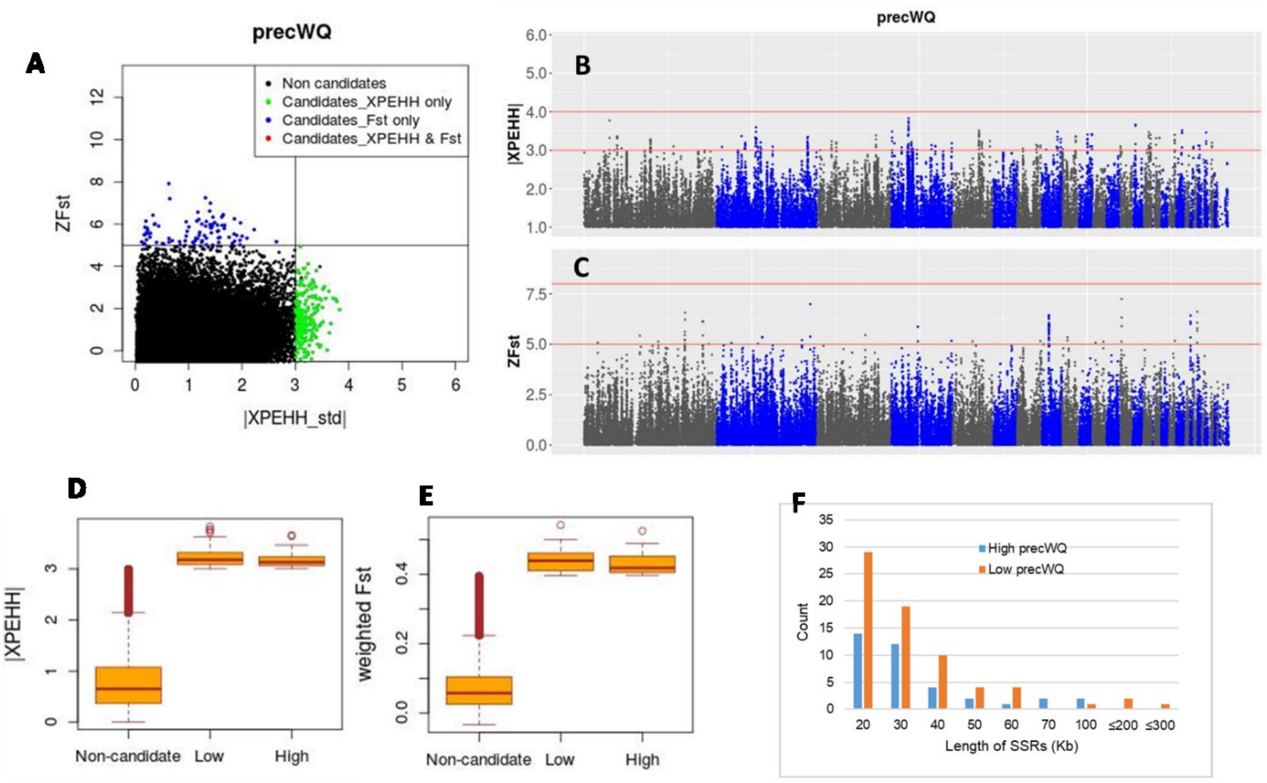


**Figure S15:** Selection signature analysis results for precWQ. (A) Scatter plot of standardaized values of XPEHH vs *Fst*. (B-C) Manhattan plots for the XPEHH and *Fst* analyses. (D-E) Box plots showing the distribution of *Fst* and XPEHH metrics for non-candidate and candidate windows from Low and High groups. (F) Length distribution of Selective Sweep Regions (SSRs).

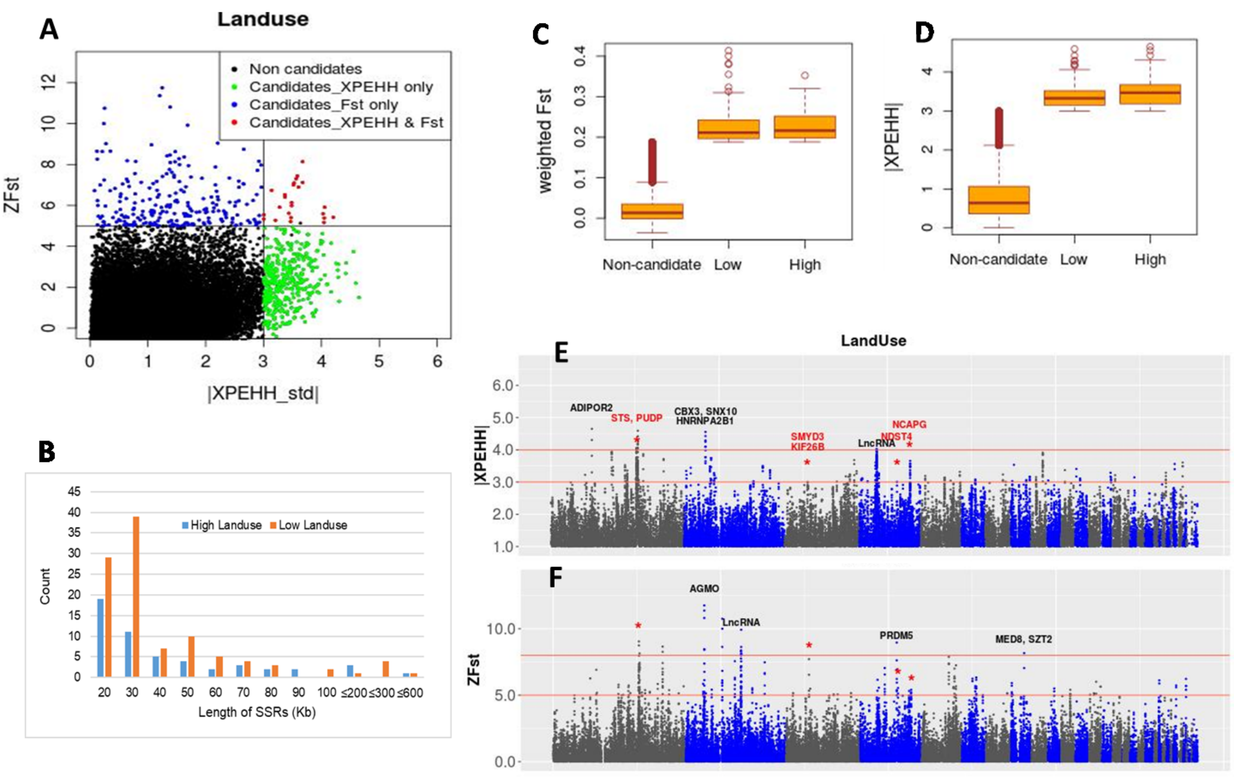


**Figure S16:** Selection signature analysis  results for Landuse.  (A) Scatter plot of standardaized values of XPEHH vs *Fst*. (B) Length distribution of Selective Sweep Regions (SSRs). (C-D) Box plots showing the distribution of *Fst* and XPEHH metrics for non-candidate windows and the candidate windows for Low and High groups. (E-F) Manhattan plots for the XPEHH and *Fst* analyses; common windows are marked with asterisk and gene names from common windows are shown in red.

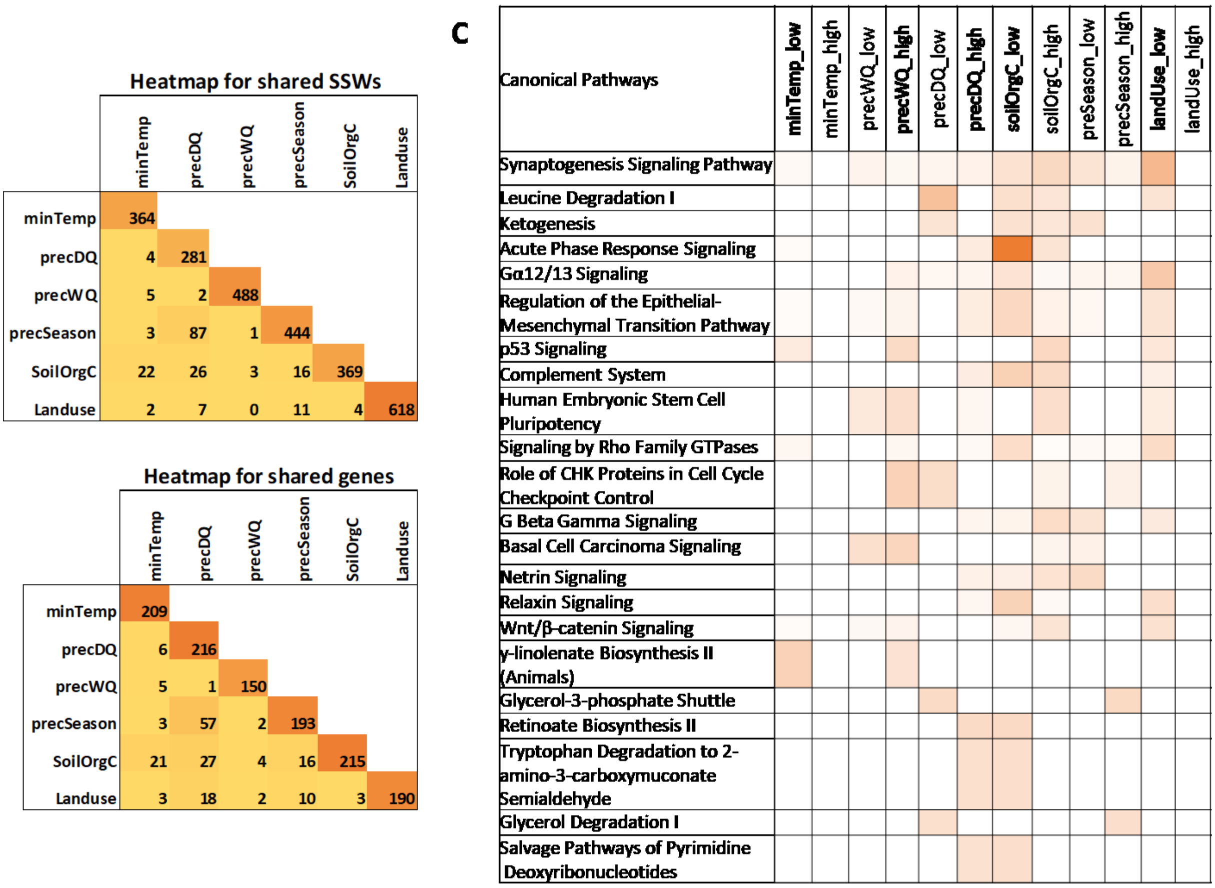


**Figure S17:** Shared selective sweep candidates among different environmental analysis. (A) Shared candidate windows; (B) shared candidate genes; (C) shared canonical pathways affected by candidate genes from different environmental analyses. Only pathways showing –log(*P*-value) ≥ 2 in at least two environmental analyses are shown.

Supplementary Tables

**Table S1. Ethiopian chicken populations studied along with the characteristics and classification of their geographic locations**

| Population | N | Agro-ecological Zone (AEZ-16) | Traditional classification | Geographic Region | District | Elevation | Slope |
|---|---|---|---|---|---|---|---|
| Adane | 10 | Tropic – cool/semiarid, sub-humid | Dega | Amhara | Kalu | 2280 - 2734 | East |
| Alfa Midir | 10 | Tropic – cool/sub-humid | Dega | Amhara | Menz Gera Midir | 3304 - 3451 | East |
| Amesha Shinkuri | 10 | Tropic – cool/sub-humid | Dega | Amhara | Fagita Lekoma | 2417 - 2564 | West |
| Arabo | 10 | Tropic – cool/semiarid | Weina - dega | Amhara | Kalu | 1624 - 1471 | East |
| Ashuda | 10 | Tropic – cool/sub-humid | Weina - dega | Amhara | South Achefer | 2009 - 2103 | West |
| Batambie | 8 | Tropic – cool/sub-humid | Weina - dega | Amhara | Banja | 2003 - 2127 | West |
| Bekele Girissa | 10 | Tropic – cool/sub-humid | Weina - dega | Oromia | Dugda | 1642 - 1668 | Center |
| Dikuli | 10 | Tropic – cool/sub-humid | Weina - dega | Amhara | South Achefer | 2022 - 2109 | West |
| Gafera | 10 | Tropic – cool/sub-humid | Dega | Amhara | Fagita Lekoma | 2441 - 2592 | West |
| Gesses | 9 | Tropic – warm/humid | Kolla | Gumuz | Dibate | 1196 - 1546 | West |
| Gijet | 9 | Tropic – cool/semiarid | Weina - dega | Tigray | Sahareti Samire | 1990 - 2303 | East |
| Hugub | 10 | Tropic – warm/semiarid | Kolla | Afar | Dulecha | 737 - 979 | East |
| Hadush Adi | 9 | Tropic – cool/semiarid | Kolla | Tigray | Merebleke | 1417 - 1669 | East |
| Jarso | 14 | Tropic – cool/semiarid | Weina - dega | Oromia | Jarso | 1817 - 2136 | East |
| Kido | 9 | Tropic – cool/ sub-humid | Kolla | Gumuz | Dibate | 1281 - 1418 | West |
| Kumato | 10 | Tropic – cool/sub-humid | Weina - dega | SNNPR | Dara | 1838 - 1969 | East |
| Local Horro | 6 | Tropic – cool/sub-humid | Weina - dega | Oromia | Horro | 1921 - 2375 | West |
| Loya | 10 | Tropic – cool/sub-humid | Weina - dega | SNNPR | Dara | 1601 - 1847 | East |
| Meseret | 10 | Tropic – cool/semiarid | Weina - dega | Tigray | Enderta | 2248 - 2312 | East |
| Metkilimat | 10 | Tropic – cool/semiarid | Weina - dega | Tigray | Sahareti Samire | 1709 - 2051 | East |
| Mihquan | 10 | Tropic – cool/semiarid | Kolla | Tigray | Merebleke | 1295 - 1495 | East |
| Negasi Amba | 10 | Tropic – cool/sub-humid | Dega | Amhara | Menz Gera Midir | 2978 - 3071 | East |
| Shubi Gemo | 10 | Tropic – cool/sub-humid | Weina - dega | Oromia | Dugda | 1643 - 1651 | Center |
| Surta | 9 | Tropic – cool/sub-humid | Dega | Amhara | Banja | 2480 - 2575 | West |
| Tsion Teguaz | 10 | Tropic – cool/semiarid | Weina - dega | Amhara | Gondar Zuria | 1870 - 1968 | West |

**Table S2. Climatic and agro-ecological variables tested in Ecological Niche Modelling for Ethiopian chicken**

| | Variable | Description | Units | Database |
|---|---|---|---|---|
| Climatic variables | Bio1 | Annual Mean Temperature | °C | WorldClim - Global climate data http://worldclim.org |
| | Bio2 | Mean Diurnal Range | °C (Bio2/Bio7) | |
| | Bio3 | Isothermality | °C | |
| | Bio4 | Temperature Seasonality | °C | |
| | Bio5 | Max temperature of warmest month | °C | |
| | Bio6 | Min temperature of coldest month | °C | |
| | Bio7 | Temperature annual range | °C (Bio5 - Bio6) | |
| | Bio8 | Mean temperature of wettest quarter | °C | |
| | Bio9 | Mean temperature of driest quarter | °C | |
| | Bio10 | Mean temperature of warmest quarter | °C | |
| | Bio11 | Mean temperature of coldest quarter | °C | |
| | Bio12 | Annual precipitation | mm/m2 | |
| | Bio13 | Precipitation of wettest month | mm/m2 | |
| | Bio14 | precipitation of driest month | mm/m2 | |
| | Bio15 | Precipitation seasonality | mm/m2 (Coeficient of variation) | |
| | Bio16 | Precipitation of wettest quarter | mm/m2 | |
| | Bio17 | Precipitation of driest quarter | mm/m2 | |
| | Bio18 | Precipitation of warmest quarter | mm/m2 | |
| | Bio19 | Precipitation of coldest quarter | mm/m2 | |
| | WatVapPress01 | Water vapour pressure of the wettest month | kPa | |
| | WatVapPress08 | Water vapour pressure of the driest month | kPa | |
| | Elevation | Meters above sea level | m.a.s.l. | |
| Soil variables | soil_pH | Soil pH | pH (x10 in H2O) | Global gridded soil information https://soilgrids.org |
| | CatEx_Capacity | Cation Exchange capacity | cmolc/Kg at depht 0.00 m | |
| | BulkD | Bulk Density | Kg/m3 at depht 0.00 m | |
| | Organic_Carbon | Soil organic carbon content | g/Kg at depht 0.00 m | |
| | Clay | Clay content | mass fraction in % at depht 0.00 m | |
| | Silt | Silt content | mass fraction in % at depht 0.00 m | |
| | Sand | Sand content | mass fraction in % at depht 0.00 m | |
| | WaterCapacity | Total available water capacity | mm2/1mt soil depht | Spatial Data Access Tool (SDAT) - NASA https://webmap.ornl.gov/ogc/wcsdown.jsp?dg_id= |
| Vegetation variables | Forest | Forest cover | % | Harmonized World Soil Dataset http://www.fao.org/soils-portal/soil-survey/soil-maps-and-databases/harmonized-world-soil-database-v12/en/ |
| | Grassland | Grass/shrub cover | % | |
| | Landuse | Land use for agricultural purposes | % | |
| | Crop_dominance | Crop dominance (majors crops) | Category | Global Food S3ecurity Analysis-Support DATA https://www.croplands.org/ |

**Table S3: Candidate selection signature regions and overlapping genes detected from analyses of different environmental parameters.**

**See File "SI_TableS3_SSRs_genes.xlsx"**

**Table S4: Candidate SNP list from selection signature analysis**

**See File "SI_TableS4_candidateSNPs_selectionSignature.xlsx".**

**Table S5: Overlap of QTLs with genes from selection signature regions.**

**See File "SI_TableS5_overlappingQTLs_SSR_genes.xlsx"**

**Table S6: Candidate SNPs and genes from RDA analysis**

**See File "SI_TableS6_candidateSNPs_RDA.xlsx"**

**Table S7A: Top molecular and cellular functions and physiological processes affected by candidate genes involved in adaptation to temperature and altitude**

| Low temperature/High altitude | High temperature/Low altitude |
|---|---|
| **<u>Top molecular and cellular function</u>**<br>- Lipid Metabolism<br>- Small Molecule Biochemistry<br>- Molecular Transport<br>- Cellular Assembly and Organization<br>- Cell-to-cell Signalling and Interactions | **<u>Top molecular and cellular function</u>**<br>- Cell Cycle<br>- Cell-To-Cell Signalling and Interaction<br>- Cellular Development<br>- Cellular Growth and Proliferation<br>- Cellular Movement |
| **<u>Physiological system development and function</u>**<br>- Renal and Urological System Development and Function<br>- Haematological System Development and Function<br>- Tissue Development<br>- Connective Tissue Development and Function<br>- Embryonic Development | **<u>Physiological system development and function</u>**<br>- Reproductive System Development and Function<br>- Nervous System Development and Function<br>- Cardiovascular System Development and Function<br>- Organ Morphology<br>- Organismal Development |

**Table S7B: Top molecular and cellular functions and physiological processes affected by candidate genes involved in adaptation to Low and High precipitation in the driest quarter (precDQ)**

| Low precipitation in driest quarter | High precipitation in driest quarter |
|---|---|
| **Top molecular and cellular function**<br>- Gene Expression<br>- Cell Morphology<br>- Cellular Assembly and Organization<br>- Cellular Movement<br>- Cellular Function and Maintenance | **Top molecular and cellular function**<br>- Cellular Assembly and Organization<br>- Cellular Function and Maintenance<br>- Molecular Transport<br>- Protein Trafficking<br>- Cell-To-Cell Signalling and Interaction |
| **Physiological system development and function**<br>- Organismal Development<br>- Nervous System Development and Function<br>- Organ Morphology<br>- Tissue Morphology<br>- Embryonic Development | **Physiological system development and function**<br>- Embryonic Development<br>- Organismal Development<br>- Tissue Morphology<br>- Nervous System Development and Function<br>- Organ Morphology |

**Table S7C: Top molecular and cellular functions and physiological processes affected by candidate genes involved in adaptation to Low and High precipitation seasonality (precSeasonality)**

| Low precipitation seasonality | High precipitation seasonality |
|---|---|
| **Top molecular and cellular function**<br>- Molecular Transport<br>- Cell Cycle<br>- Cell-To-Cell Signalling and Interaction<br>- Cellular Assembly and Organization<br>- Cellular Development | **Top molecular and cellular function**<br>- Cell Death and Survival<br>- Gene Expression<br>- Cell Cycle<br>- Cellular Movement<br>- Cell Morphology |
| **Physiological system development and function**<br>- Connective Tissue Development and Function<br>- Embryonic Development<br>- Skeletal and Muscular System Development and Function<br>- Tissue Development | **Physiological system development and function**<br>- Cardiovascular System Development and Function<br>- Organ Morphology<br>- Organismal Development<br>- Renal and Urological System Development and Function<br>- Embryonic Development |

**Table S7D: Top molecular and cellular functions and physiological processes affected by candidate genes involved in adaptation to Low and High precipitation in the wettest quarter (precWQ)**

<table>
<tr><th>Low precipitation in wettest quarter</th><th>High precipitation in wettest quarter</th></tr>
<tr><td><u>Top molecular and cellular function</u><br>- Amino Acid Metabolism<br>- Carbohydrate Metabolism<br>- Cell Death and Survival<br>- Cell Morphology<br>- Cell-To-Cell Signalling and Interaction</td><td><u>Top molecular and cellular function</u><br>- Cellular Assembly and Organization<br>- Cellular Function and Maintenance<br>- Cell Cycle<br>- DNA Replication, Recombination, and Repair<br>- Cellular Development</td></tr>
<tr><td><u>Physiological system development and function</u><br>- Embryonic Development<br>- Organismal Development<br>- Tissue Development<br>- Organ Development<br>- Respiratory System Development and Function</td><td><u>Physiological system development and function</u><br>- Nervous System Development and Function<br>- Tissue Morphology<br>- Cardiovascular System Development and Function<br>- Embryonic Development<br>- Organ Development</td></tr>
</table>

**Table S7E: Top molecular and cellular functions and physiological processes affected by candidate genes involved in adaptation to Low and High soil organic carbon content (SoilOrgC)**

| Low soil organic carbon content | High soil organic carbon content |
|---|---|
| **Top molecular and cellular function**<br>- Cellular Function and Maintenance<br>- Molecular Transport<br>- Cellular Assembly and Organization<br>- Cell-To-Cell Signalling and Interaction<br>- Cell Death and Survival | **Top molecular and cellular function**<br>- Cell-To-Cell Signalling and Interaction<br>- Cellular Assembly and Organization<br>- Cellular Function and Maintenance<br>- Cellular Development<br>- Cellular Growth and Proliferation |
| **Physiological system development and function**<br>- Hematological System Development and Function<br>- Tissue Development<br>- Renal and Urological System Development and Function<br>- Organismal Functions<br>- Tissue Morphology | **Physiological system development and function**<br>- Nervous System Development and Function<br>- Tissue Development<br>- Cardiovascular System Development and Function<br>- Organ Morphology<br>- Organismal development |

**Table S7F: Top molecular and cellular functions and physiological processes affected by candidate genes involved in adaptation to Low and High proportion of land use for cultivation purpose (LandUse)**

| Low proportion of land used for cultivation | High proportion of land used for cultivation |
|---|---|
| **Top molecular and cellular function**<br>- Nucleic Acid Metabolism<br>- Small Molecule Biochemistry<br>- Molecular Transport<br>- Cellular Movement<br>- Drug Metabolism | **Top molecular and cellular function**<br>- Cell-To-Cell Signalling and Interaction<br>- Cellular Assembly and Organization<br>- Amino Acid Metabolism<br>- Lipid Metabolism<br>- Molecular Transport |
| **Physiological system development and function**<br>- Embryonic Development<br>- Organismal Development<br>- Nervous System Development and Function<br>- Organ Development<br>- Tissue Development | **Physiological system development and function**<br>- Behaviour<br>- Embryonic Development<br>- Nervous System Development and Function<br>- Organismal Development Tissue Development |

**Table S8: Important candidate genes associated with different environmental predictors from RDA analysis. The list include those genes associated with SNPs with r-value ≥ 0.4 with an environmental predictor or are common between at least one SSA approach and RDA analysis with environmental r value>=0.3 (shown with *).**

| Candidate genes | Associated SNP, type & correlation | Gene function |
|---|---|---|
| ***minTemp*** | | |
| *ULK3* | chr10_2871082 (intronic); r=0.42 | Acts as a regulator of autophagy following cellular senescence (cell aging process stimulated in response to cellular stress) (1) |
| *PITX2* | chr4_57798697 (UTR3); *r*=0.40 | Acts as transcription factor; critical in heart development (1); promotes heart cell repair in response to ischemic injury (2) |
| *CDH8* | chr11_12943908 (intronic); *r*=0.40 | Involved in response to cold (1) |
| *UTP18** | chr18_5139670 (Intergenic); *r*=0.36 | rRNA processing (1) |
| *FSTL4** | chr13_17178369 (intergenic); *r*=0.34 | Possibly involved in stress-induced cardiac hypertrophy (3) |
| *ENSGALG 00000052790** | chr1_37163689 (ncRNA_intronic); r=0.33 | LncRNA with possible cis-regulatory role on nearby protein-coding genes. |
| *CA10* | chr18_5158248 (Intronic); r=0.32 | Brain development (1) |
| *ENSGALG 00000042572 (TIMM10B)** | chr1_197440205 (Downstream); r=0.31 | *TIMM10B* is a component of the TIM22 complex that mediates the import and insertion of multi-pass transmembrane proteins into the mitochondrial inner membrane (1). |
| ***precDQ*** | | |
| *PRR5L* | chr5_19713610 (Intergenic), *r*=0.44 | Cellular response to oxidative stress (1) |
| *ENSGALG 00000052510* | chr2_7014078 (ncRNA_intronic), *r*=0.41 | LncRNA with possible cis-regulatory role on nearby protein-coding genes. |
| ***precWQ*** | | |
| *ENSGALG 00000007080* and *GDF9* | chr13_17762839 (Intergenic); r=0.50 | *ENSGALG00000007080:* Novel protein coding gene<br>GDF9 is a growth differentiation factor, important in oocyte growth and development (1). |
| *CPQ* and *ENSGALG 00000048883* | chr2_127127333 (intergenic); r=0.49 | CPQ have important roles in proteolysis, and possibly involved in thyroid hormone generation, and tissue regeneration (1)<br>ENSGALG00000048883 : function unknown |
| ENSGALG 00000044225 (Zinc finger protein 132-like) | chr4_70549769 (Intronic); r= 0.48 | Zink Finger proteins (ZNFs) are involved in the regulation of several cellular processes. In fact, ZNFs are implicated in transcriptional regulation, ubiquitin-mediated protein degradation, signal transduction, actin targeting, DNA repair, cell migration, and numerous other processes (4). |
| *ZNF800* | chr1_21037106 (downstream); *r*=0.48 | ZNFs are implicated in transcriptional regulation, ubiquitin-mediated protein degradation, signal transduction, actin targeting, DNA repair, cell migration, and numerous other processes (4). |
| *DCT* and *ENSGALG* | chr1_147942295 (intergenic); r=0.45 | DCT is involved in melanin biosynthesis pathway (1) |

| | | |
|---|---|---|
| *00000016900* *(*Glypican 6 or GPC6) | | The glypican encoded by this gene is a putative cell surface coreceptor for growth factors, extracellular matrix proteins, proteases and anti-proteases (1) |
| *FOLR1* and *ENSGALG 00000017342 (OR52B2L)* | chr1_197330375 (Intergnic); r=0.45 | FOLR1: Required for normal embryonic development and normal cell proliferation.<br>OR52B2L: Olfactory receptor activity (1) |
| *LRRC7* | chr8_29320541 (Intronic); *r*=0.45 | Involved in nervous system processes (1) |
| *TMEM178B* | chr1_57151740 (Intronic), *r*=0.45 | A transmembrane protein gene (1) |
| *SUPT3H* | chr3_109090270 (Intronic), *r*=0.45 | Transcription initiation (1) |
| *RELL1* | chr4_69847804 (Intronic), *r*=0.45 | Induces apoptosis, when overexpressed |
| *LCP2* | chr13_4628240 (Intronic); *r*=0.43 | Immune response (1) |
| *FLOT2* | chr19_6023827 (Intronic); *r*=0.42 | Epidermal cell adhesion and epidermis development and function; neural signalling (1) |
| *PACRGL* | chr4_74948889 (Intronic); r=0.42 | Function yet unknown |
| *EXOC1L* and *NMU* | chr4_65022119 (intergenic); r=0.42 | EXOC1L: Golgi to plasma membrane transport<br>NMU: Stimulates muscle contractions of specific regions of the gastrointestinal tract. Affects many biological processes including eating behaviour, energy homeostasis, gastric acid secretion, regulation of heart rate, sensory perception of pain, temperature homeostasis (1) |
| *LMO1* | chr5_10353784 (intronic) ; r=0.41 | May be involved in gene regulation within neural lineage cells potentially by direct DNA binding or by binding to other transcription factors. (1) |
| *ENSGALG 00000052884* | chr1_44464131 (ncRNA_intronic); r=0.41 | LncRNA with possible cis-regulatory role on nearby protein-coding genes. |
| *KIF2B and ENSGALG 00000048193* | chr18_5660027 (intergenic) | *KIF2B*: regulation of chromosome segregation (1)<br>*ENSGALG00000048193: LncRNA)* |
| *ENSGALG 00000051497* and *ERICH1* | chr3_91444645 (intergenic); r=0.40 | LncRNA with possible cis-regulatory role on nearby protein-coding genes.<br>*ERICH1*: function unknown |
| *ENSGALG 00000033453* and *FAM49B* | chr2_140333337 (Intergenic); r=0.40 | LncRNA with possible cis-regulatory role.<br>*FAM49B*: regulator of mitochondrial function and integrity that suppresses tumor metastasis (5) |
| *TEKT3* | chr18_2306225 (Intronic); r=0.40 | May be a structural component of the sperm flagellum. Required for normal sperm mobility (1) |
| *SUSD4* | chr3_17861357 (Intronic); *r*=0.40 | Innate immune response (1) |
| *ADAM33** | chr4_89589528 (Intronic); *r*=0.40 | Involved in proteolysis, which is an important adaptive mechanism in plants in response to severe water deficit condition (6) |
| *RASSF3* | chr1_33839618 (Intronic); *r*=0.39 | Involved in apoptotic process and signal transduction (1) |
| *OTUD7B* | chr25_1809778 (Intronic); *r*=0.39 | Adaptive immune response (1) |
| *ARHGEF17* | chr1_196484159 (Intronic); *r*=0.39 | Involved in apoptotic process and signal transduction (1) |

| | | |
|---|---|---|
| *PLA2G4A** | chr8_10641466 (Intronic); *r*=0.36 | Major role in membrane lipid remodelling and biosynthesis of lipid mediators of the inflammatory response (1) |
| *SLC22A23** | chr2_66254341 (Intronic); *r*=0.35 | Transmembrane transporter activity (1) |
| *PAX3** | chr9_8044315 (Intronic); *r*=0.32 | Transcription factor regulating cell proliferation, migration, and apoptosis. Involved in neural development and myogenesis (1) |
| ***precSeasonality*** | | |
| *ST3GAL6* | chr1_85341100 (intronic), *r*=0.46 | Involved in cellular response to Interleukin-6 which plays important role in oxidative stress response (1) |
| *WIF1* | chr1_34033189 (Intronic), *r*=0.43 | Regulation of Wnt signalling pathway, which negatively regulates stress granule assembly in cells (7) |
| *ENSGALG 00000046974* | chr1_172280958 (intergenic), *r*=0.42 | LncRNA with possible cis-regulatory function. |
| *HTR2C** | chr1_85341100 (Intronic), *r=0.34* | Crucial mediatory role in the stress-induced activation of hypothalamic-pituitary-adrenal (HPA) axis (8). |
| *SLIT3** | chr13_5424064 (Intronic); r=0.33 | Many functions including apoptotic process, atrioventricular valve morphogenesis, axon guidance, cellular response to hormone stimulus, negative chemotaxis, negative regulation of cell growth, negative regulation of cell population proliferation, response to cortisol, Roundabout signalling pathway (1) |
| *EPHA7** | chr3_73992880 (Intronic), *r=0.32* | Involved in many biological processes (See Table 1) |
| ***SoilOrgC*** | | |
| *KIF5C* | chr7_34800723 (Intronic), *r*=0.50 | Involved in synaptic transmission, axonal protein transport, axon guidance (1) |
| *ZEB2** | chr7_33312018 (Intronic), *r*=0.47 | Many important functions in brain and nervous system development and processes. Other roles include positive regulation of lens fiber cell differentiation, positive regulation of melanin biosynthetic process, and melanocyte differentiation (1) |
| *RAD54B* | chr2_126027584 (Upstream), *r*=0.47 | Involved in DNA repair and mitotic recombination(1); found upregulated in the hypothalamus of fast growing chicken line (9) |
| *ZNF536* | chr11_8674383 (intergenic), *r*=0.43 | Involved in transcriptional regulation including regulation of neuron differentiation (1) |
| *URI1* | chr11_8674383 (intergenic), *r*=0.43 | Involved in cellular response to growth factor, cellular response to steroid hormone stimuli, and regulation of cell growth (1) |
| *ENSGALG 00000035994* | chr2_23318735 (intronic); r=0.43 | Function unknown |
| *ENSGALG 00000036131* | chr1_52325047 (Intergenic); r=0.42 | LncRNA |
| *ENSGALG 00000048545** | chr2_9781217 (Intergenic); r=0.38 | LncRNA |
| *CDH2** | chr2_104898963 (Intronic); r=0.38 | Encodes a cell adhesion protein with many functions in brain and neuronal development and nervous-system processes (1) |
| *FOXO3** | chr3_67330792 (intronic), *r*=0.36 | Regulates many functions e.g. aging, brain morphogenesis, cellular response to nerve growth factor stimulus, cellular response to glucose stimulus, oxidative stress and DNA damage responses, positive regulation of apoptosis and autophagy in response to starvation (1) |
| *FARP1** | chr1_146492072 (Intronic), *r*=0.36 | Involved in nervous system processes such as dendrite morphogenesis and synapse assembly (1) |
| *NADSYN1** | chr5_1889541 (Intronic); r=0.35 | Involved in Nicotinamide adenine dinucleotide (NAD) biosynthetic and metabolic processes (1). NAD is s a coenzyme in metabolic redox |

| | | |
|---|---|---|
| | | reactions, a precursor for several cell signaling molecules, and a substrate for protein posttranslational modifications (10) |
| *C8orf37** | chr2_126570145 (Intergenic); r=0.34 | Photoreceptor cell morphogenesis (1) |
| *ADGRL3* | chr4_48035555 (Intronic); r=0.30 | Plays a role in cell-cell adhesion and neuron guidance. Important in determining the connectivity rates between the principal neurons in the cortex. (1) |
| ***LandUse*** | | |
| *ENSGALG 00000019300 (TH-like gene)* | chr1_55126703 (Intronic); *r*=0.43 | *TH* gene is involved in many functions including regulation of eating behaviour, response to nutrient level, response to growth hormone, epinephrine biosynthetic process, and eye photoreceptor cell development (1) |
| *ENSGALG 00000040732* | chr4_11524544 (ncRNA_intronic); *r*=0.43 | LncRNA, possible regulatory function |
| *WASHC3** | chr1_55453148 (Intronic); r=0.42 | Actin filament polymerization, exocytosis, protein transport (1) |
| *ENSGALG 00000048732* | chr1_54963708 (Intergenic); r=0.42 | LncRNA with possible cis-regulatory function |
